# Water Stress and Disruption of Mycorrhizae Induce Parallel Shifts in Phyllosphere Microbiome Composition

**DOI:** 10.1101/2021.04.29.442000

**Authors:** Reena Debray, Yvonne Socolar, Griffin Kaulbach, Aidee Guzman, Catherine A. Hernandez, Rose Curley, Alexander Dhond, Timothy Bowles, Britt Koskella

## Abstract

Water and nutrient limitation are key stressors that affect plant health and ecosystem function. These environmental factors impact both soil- and root-associated microbial communities, and systemically alter plant physiology, but it is less clear how they affect aboveground plant-microbe interactions. Through experimental manipulations in the field and growth chamber, we examined the interacting effects of irrigation, soil fertility, and root mycorrhizal associations on bacterial and fungal communities of the tomato phyllosphere (*Solanum lycopersicum*). Both water stress and mycorrhizal disruption reduced bacterial richness within plants, homogenized bacterial community diversity among plants, and reduced the relative abundance of dominant fungal taxa. We observed striking parallelism in the individual microbial taxa affected by irrigation and mycorrhizal associations. Given the increasingly understood role of the phyllosphere in shaping plant health and pathogen susceptibility, these results offer an additional mechanism by which belowground conditions shape plant fitness.

## INTRODUCTION

Soil water and nutrient content are among the most important predictors of plant health and biodiversity (Pautasso et al. 2010; Gupta, Rico-Medina, and Caño-Delgado 2020). Disturbances or regime shifts can deplete water and nutrients from soil, and are expected to increase in frequency and severity in the coming years (Seidl et al. 2017; Pritchard 2011; St.Clair, St. Clair, and Lynch 2010). It is thus crucial to understand how diverse and interacting stressors act to shape plant function. An important piece of the puzzle is the impacts of these changes on plant-associated microbial communities. Under stressful conditions, such as drought or pathogen infection, plants can secrete root exudates that enrich for specific microorganisms that improve plant stress tolerance (Xu et al. 2018; Carrión et al. 2019). Microbial communities associated with plant foliage, known as the phyllosphere, are increasingly recognized in their influence on plant health, including through nutrient fixation, tolerance of extreme temperatures, and protection against pathogens (Berg and Koskella 2018; Fürnkranz et al. 2008; Hubbard, Germida, and Vujanovic 2014; Lindow, Arny, and Upper 1982). Yet, much less is known about how soil properties alter phyllosphere microbiome composition or whether such changes can be adaptive for host plants.

To capture effects of soil conditions on the phyllosphere microbiome, we focused on several soil properties that induce or modulate systemic responses in plant physiology. Water stress can impact nutrient and biomass allocation, phytohormone production, and toxin accumulation in leaves (Zhou et al. 2019; Moles et al. 2018; Gupta, Rico-Medina, and Caño-Delgado 2020), while soil fertility impacts nutrient content and acidity of aboveground plant tissues (Fageria and Barbosa Filho 2007; Wang, Li, and Malhi 2008). Importantly, while soil fertility commonly improves plant reproductive output, it can also increase pest and pathogen sensitivity, potentially through alterations of below-ground microbial associations (Johnson 1993; Weese et al. 2015; Berg and Koskella 2018). We therefore also manipulated a biotic interaction that plays a major role in modulating plant responses to water and nutrient limitation. Arbuscular mycorrhizal fungi (AMF) colonize plant roots and induce systemic changes in shoots, including regulation of defense- and stress-associated genes (Gerlach et al. 2015; J. Liu et al. 2007). Like other components of soil health, plant-mycorrhizal symbioses are increasingly vulnerable to disruption under climate and land use changes (Rillig, Treseder, and Allen 2002; Verbruggen and Toby Kiers 2010).

In this study, we first asked how water stress, phosphate fertility, and mycorrhizal colonization affected bacterial and fungal communities in the tomato plant phyllosphere (*Solanum lycopersicum*). We expected that water stress would decrease microbial biodiversity, but that mycorrhizae, which buffer drought stress in plants, would minimize the effects of water stress on aboveground microbiota. In contrast, because nutrient-supplemented plants reduce carbon allocations to mycorrhizae (Johnson 1993), we predicted that organic fertilization would counteract effects of mycorrhizae on phyllosphere microbiome composition. We then asked whether stress-induced changes in microbiome composition can be adaptive by transplanting communities from a subset of treatments onto juvenile tomato plants in the growth chamber. We expected that plants grown under mycorrhizae- or nutrient-limited conditions in the growth chamber would have increased performance (as measured by resistance to the bacterial pathogen *Pseudomonas syringae*) if they received a microbial inocula from mycorrhizae- or nutrient-limited conditions in the field, compared to inocula from benevolent field conditions. This combination of field and growth chamber studies allowed us to test how soil conditions shape the diversity, composition, and function of bacteria and fungi in the phyllosphere. We found that water stress and mycorrhizal colonization were the strongest predictors of both bacterial and fungal composition. Contrary to our expectations, these factors did not interact, and in fact were independently associated with parallel shifts in microbial community composition.

## METHODS

### Field site and treatments

The field trial was conducted at the University of California Davis Student Farm (38°32′29.49″N, 121°46′0.94″W), a certified organic farm, during the 2019 growing season. Three treatments, water stress, phosphorus fertility, and mycorrhizal associations, were arranged in a split plot design and assigned to levels in a P fertility gradient to create an experimental regression (**Figure S1**). Water stress included two levels, fully watered (100% ET_c_) and deficit (50% ET_c_) irrigation applied in alternating beds, with the deficit beginning 22 days after transplant, and water completely cut to the deficit beds 76 days after transplant. Phosphorus fertility was manipulated as a 12-level gradient of phosphorus established in each irrigation condition (24 plots total) to establish an experimental regression. We applied an organic phosphorus source, seabird guano (0-11-0, Down to Earth) on the surface of each plot, which was then incorporated with a cultivator a week before planting. The gradient ranged from 0-550 grams of P (0-5000g fertilizer) per square meter of soil, including duplicate plots of the lowest (0 g P/m^2^) and highest (550 g P/m^2^) treatments **(Table S1)**. The 24 experimental plots were each split into two subplots, one with a tomato genotype deficient in mycorrhizal associations (*reduced mycorrhizal colonization*, (Barker et al. 1998)) and another with its wild-type progenitor (76R), for a total of 48 experimental units. 76R and *rmc* seeds were sourced from plants grown in the greenhouse. To avoid effects of adjacent irrigation treatments, 3 buffer plants were planted but not sampled between experimental units.

### Sample collection from field

Two months after transplant, 3 compound leaves per plant were sampled from both irrigation regimes and genotypes along the two lowest (0 g P/m^2^), two middle (220 and 330 g P/m^2^), and two highest (550 g P/m^2^) positions in the phosphorus gradient. Four plants were sampled per replicate plot, for a total of 8 plants per combination of irrigation regime, plant genotype, and phosphorus treatment. Samples were transported to the lab on ice. Samples used to generate inocula for growth chamber experiments were briefly stored at 4°C before processing, while samples used only for sequencing were frozen at –20°C.

To isolate microbial communities, leaves were pooled from the same plant into bags and suspended in sterile buffer. Bags were submerged in a Branson M5800 sonicating water bath for 10 minutes to gently dislodge microbial cells from the leaf surface. The resulting leaf wash was centrifuged at 3500 x *g* for 10 minutes, resuspended in sterile buffer, and frozen at –80°C in a KB-glycerol cryoprotectant. Up to 16 individual plants were processed each day until complete. Plant sonication batches were randomly assigned with respect to treatments, and were included as a covariate in all statistical models.

Arbuscular mycorrhizal fungi colonization rates were determined using roots collected during the midseason sampling period (51 days after transplant). Using a 6 cm dia. × 10 cm deep core, plant roots were collected 10 cm from the stem from 1 plant per subplot. Samples were rinsed in water and stored in ethanol. Roots were bleached and stained with Parker Quink black ink. Stained roots were mounted on a microscope slide and analyzed using the gridline intersect method (Giovannetti and Mosse 1980). Plant-available phosphorus was measured at all soil sampling times (51 and 116 days after transplant) using the Olsen P method.

### Plant care in growth chamber experiments

Surface-sterilized tomato seeds were germinated on sterile agar, then transplanted into pots (See Supplemental Material for growth chamber conditions). In the mycorrhizae experiment, an inocula containing equal amounts of several arbuscular mycorrhizal fungi taxa (*Gigaspora rosea, G. albida, Acaulospora spinosa, A. spinosa, Rhizophagus intradices*, and *Funneliformis mosseae*) was added to each pot at the transplant stage. AMF inoculum was acquired from INVAM (West Virginia University, Morgantown, WV). Half of the inocula (14 grams) was mixed into the potting mix, and the other half was added directly to the seedling transfer site at the top. In the phosphorus experiment, 105.6 mg of P (960 mg fertilizer) was added to half of the pots at the transplant stage, approximating the highest position of the phosphorus gradient in the field (550 g P/m^3^, assuming a fertilizer penetration depth of 20 cm). The other half of the pots remained unfertilized (0 g P/m^3^). All pots were immediately randomized with respect to treatment and remained under controlled growth chamber conditions for the duration of the experiment.

### Microbiome spray and pathogen challenge

Three-week-old plants were sprayed with microbial inocula from tomato plant leaves in the field. To track assembly of the same source communities under different conditions, each sample (which corresponded to a single plant in the field) was divided in half and used to generate inocula for two plants. In the mycorrhizae experiment, each field sample was used to inoculate one wild-type plant and one *rmc* plant in the growth chamber. In the phosphorus experiment, each field sample was used to inoculate one fertilized and one unfertilized plant in the growth chamber. Each plant was sprayed with 4 mL of microbial inocula at a standardized concentration of 10^4^ CFU/mL.

One week after spraying with communities from the field, plants were experimentally infected with the bacterial pathogen *Pseudomonas syringae* at an optical density of 0.0002 (approximately 5 × 10^3^ CFU/mL). Three leaves per plant were inoculated by blunt-end syringe injection into the abaxial side of the leaf.

24 hours after the *P. syringae* challenge, plants were destructively sampled. Three hole punches were taken from each of the inoculated leaves and homogenized in sterile buffer using a FastPrep-24 5G sample disruption instrument. *Pseudomonas* density on leaves was obtained through CFU plating and droplet digital PCR (ddPCR). The remaining aboveground plant material, which had been sprayed with microbial inocula but not directly challenged with *P. syringae*, was collected in a 15 mL tube. Microbial communities were isolated through sonication and centrifugation as described above. Total bacterial density was obtained through ddPCR with primers for conserved sequences in the 16S rRNA gene. Primer sequences and reaction conditions for ddPCR assays are available in the Supplemental Material.

### DNA extraction and sequencing

DNA was extracted using the MoBio PowerSoil DNA isolation kit. Extraction batches of 24 samples were assigned randomly with respect to treatments and included in all statistical models. One extraction in each set of 24 was a negative control. Libraries were prepared by amplifying the V4 region of the 16S rRNA gene and the ITS2 gene (see Supplementary Material for primer sequences). Libraries were amplified, cleaned up, and sequenced alongside DNA extraction controls and PCR controls on the Illumina MiSeq platform at Microbiome Insights (Vancouver, BC, CAN).

Reads were analyzed using the recommended DADA2 workflow (Callahan et al. 2016) to infer amplicon sequencing variants. Taxonomy was assigned using the SILVA and UNITE databases (Quast et al. 2013; Kõljalg et al. 2005). Because many ITS sequences could not be classified at the kingdom level by UNITE, the hidden Markov model classifier ITSx was used to further delineate ITS sequences and remove non-fungal sequences (Bengtsson-Palme et al. 2013). 16S variants were filtered to remove chloroplast and mitochondrial sequences. Potential contaminants were filtered based on prevalences in negative controls versus true samples using the package *decontam* (Davis et al. 2018).

### Statistical analysis

Analyses of alpha and beta diversity were conducted using ANOVA and PERMANOVA tests with water regime, plant genotype, soil fertilizer concentration (g/m^2^), plant sonication batch, and DNA extraction batch as covariates. When treatments were allowed to interact, none of the interaction terms were significant, and model selection based on Akaike’s information criterion favored the model without interactions. Indicator taxa were identified based on Dufrêne and Legendre’s indicator value (R package *multipatt*). To independently validate this analysis, we also modeled differential abundance across treatments (R package *edgeR*) (Robinson, McCarthy, and Smyth 2010).

The correlation between bacterial and fungal richness were assessed with Pearson’s product correlation coefficient. Partial R^2^ values were calculated to test whether this correlation could be attributed to shared responses of bacteria and fungi to other variables (R package *rsq*). To test correlations between bacterial and fungal composition, a partial Mantel test was conducted on their respective Bray-Curtis dissimilarity matrices along with a Euclidean distance matrix of field trial treatments and technical batches. To construct a co-occurrence network, the abundances of the top 500 bacterial taxa and the top 500 fungal taxa were first converted to presence-absence measures (as relative abundances would not be directly comparable across sequencing libraries). The two datasets were combined and the package *rcorr* was used to generate matrices of Pearson’s correlations and corresponding *p-*values. After Benjamini-Hochberg correction for multiple testing (FDR<0.1), a network was constructed on significant correlations (R package *igraph*).

While colonization of plants in the growth chamber by field communities was generally high, a subset of plants had low bacterial abundance and a high proportion of ASVs unique to the growth chamber, which appeared to confound estimates of bacterial diversity (**Figure S2**). To address this, ASVs were classified as shared with the field trial or unique to the growth chamber, and all analyses were conducted separately on the two categories. In support of this approach, ASVs shared with the field trial generally recapitulated treatment effects observed in the field, while the potential growth chamber contaminants appeared to be randomly distributed with respect to treatment (**Figure S3**). A paired Wilcoxon signed-rank test was used to compare microbiome assembly on plants in the growth chamber that had received the same inoculum from the field.

## RESULTS

### Effects of experimental treatments on soil conditions

In the field, wild-type plants had significantly higher AMF root colonization than the reduced mycorrhizal genotype (mean 76R = 17%, mean *rmc* = 11%; 95% CI for difference in means = [0.60 11], p = 0.032; **Figure S4**). Though there were no significant differences in soil gravimetric water content between fully watered and deficit irrigation plots at the mid-season sampling (51 days after transplant), there was a significant decrease in soil water content at all three depths (0-15cm, 15-30cm, 30-60cm) at harvest (116 days after transplant). The applied P gradient did not significantly impact available soil P (Olsen P) at any of the sampling events.

### Phyllosphere microbiome composition in field trial

We first measured the impact of the field trial treatments on microbial community composition in the phyllosphere. Water regime and mycorrhizal associations significantly predicted variation in bacterial and fungal community composition, while the effect of fertilization was marginally significant in fungal communities only (**Table 1)**.

**Table 1.** Full results of PERMANOVA models on Bray-Curtis dissimilarity matrices.

| **Bacterial community composition** |  |  |  |  |  |  |
| --- | --- | --- | --- | --- | --- | --- |
| **Treatment** | **Df** | **Sum of squares** | **Mean square** | **F value** | **R<sup>2</sup>** | **p-value** |
| Water regime | 1 | 0.601 | 0.601 | 1.496 | 0.015 | **0.00079** |
| Genotype | 1 | 0.495 | 0.495 | 1.232 | 0.012 | **0.028** |
| Fertilizer concentration | 1 | 0.455 | 0.455 | 1.133 | 0.011 | 0.112 |
| Plant sonication batch | 12 | 5.763 | 0.480 | 1.196 | 0.145 | **<0.0001** |
| DNA extraction batch | 6 | 3.427 | 0.571 | 1.423 | 0.086 | **<0.0001** |
| **Fungal community composition** |  |  |  |  |  |  |
| **Treatment** | **Df** | **Sum of squares** | **Mean square** | **F value** | **R<sup>2</sup>** | **p-value** |
| Water regime | 1 | 0.470 | 0.470 | 1.436 | 0.014 | **0.011** |
| Genotype | 1 | 0.482 | 0.482 | 1.473 | 0.015 | **0.0089** |
| Fertilizer concentration | 1 | 0.404 | 0.404 | 1.234 | 0.013 | 0.061 |
| Plant sonication batch | 12 | 4.633 | 0.386 | 1.180 | 0.141 | **0.0003** |
| DNA extraction batch | 6 | 3.388 | 0.564 | 1.726 | 0.103 | **<0.0001** |

Shifts in community composition often reflected changes in alpha diversity (species diversity per plant) and beta diversity (heterogeneity within treatments) (full model results in **Tables S2-S3**). Compared to fully watered plants, water-stressed plants harbored lower bacterial richness (F=10.468, *p*=0.0018) and more similar communities to one another (F=7.347, *p*=0.0080), while Shannon diversity was not affected. Simultaneous decreases in alpha and beta diversity often indicate loss of rare species (i.e. species that inhabit some sites, but not others, across a landscape) (Socolar et al. 2016). Accordingly, rare ASVs showed the strongest discrepancies between the full water and deficit treatments (**Figure S5**). In fungal communities, water stress increased the Shannon diversity index (F=7.966, *p*=0.0062), a combined measure of richness and evenness. Further examination of the individual components revealed that this effect was driven by an increase in evenness under water-limited conditions (F=8.495, *p*=0.0047), while richness was not affected **(Figure 1)**.

**Figure 1.**
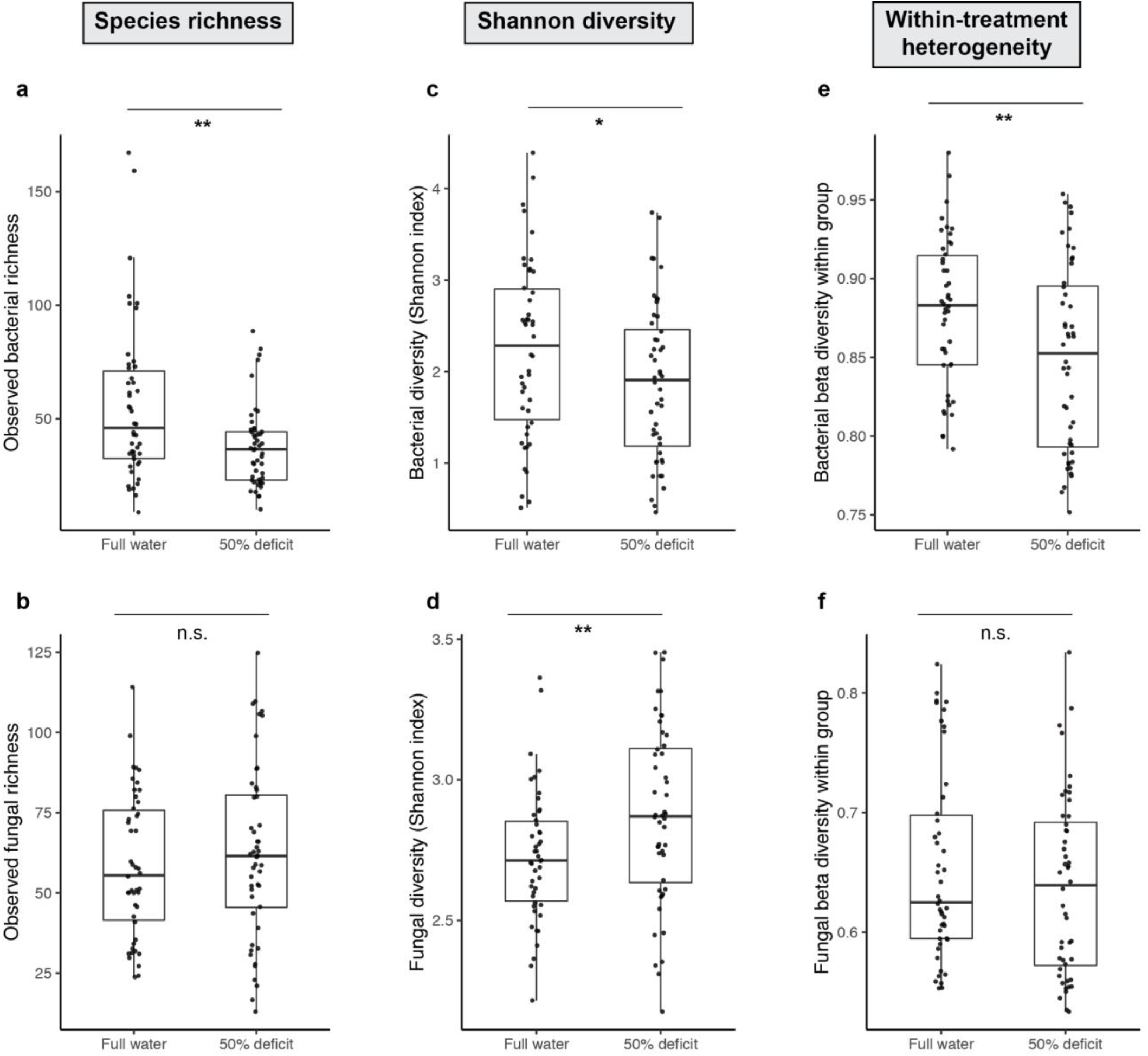
Water stress reduces bacterial alpha and beta diversity and increases fungal alpha diversity in the tomato phyllosphere. **(a**,**b)** ASV counts after removing plant sequences and contaminant sequences. Bacterial richness values were log-transformed prior to ANOVA to meet normality and homoscedasticity assumptions. **(c**,**d)** Shannon-Weaver index of combined richness and evenness. **(e**,**f)** Within-group dispersion was quantified by generating a Bray-Curtis dissimilarity matrix. Points represent the mean Bray-Curtis distance of each sample to other samples within the same treatment group (full water or 50% deficit).

The reduced mycorrhizal (*rmc*) plants harbored lower phyllosphere bacterial richness (F=5.203, *p*=0.026) and more similar communities to one another (F=7.548, *p*=0.0072) than their wild-type counterparts. As with water stress, the strongest discrepancies between wild-type and *rmc* plants were among rare ASVs (**Figure S6**). The *rmc* plants harbored lower fungal richness (F=11.417, *p*=0.0011) than wild-type plants **(Figure 2)**.

**Figure 2.**
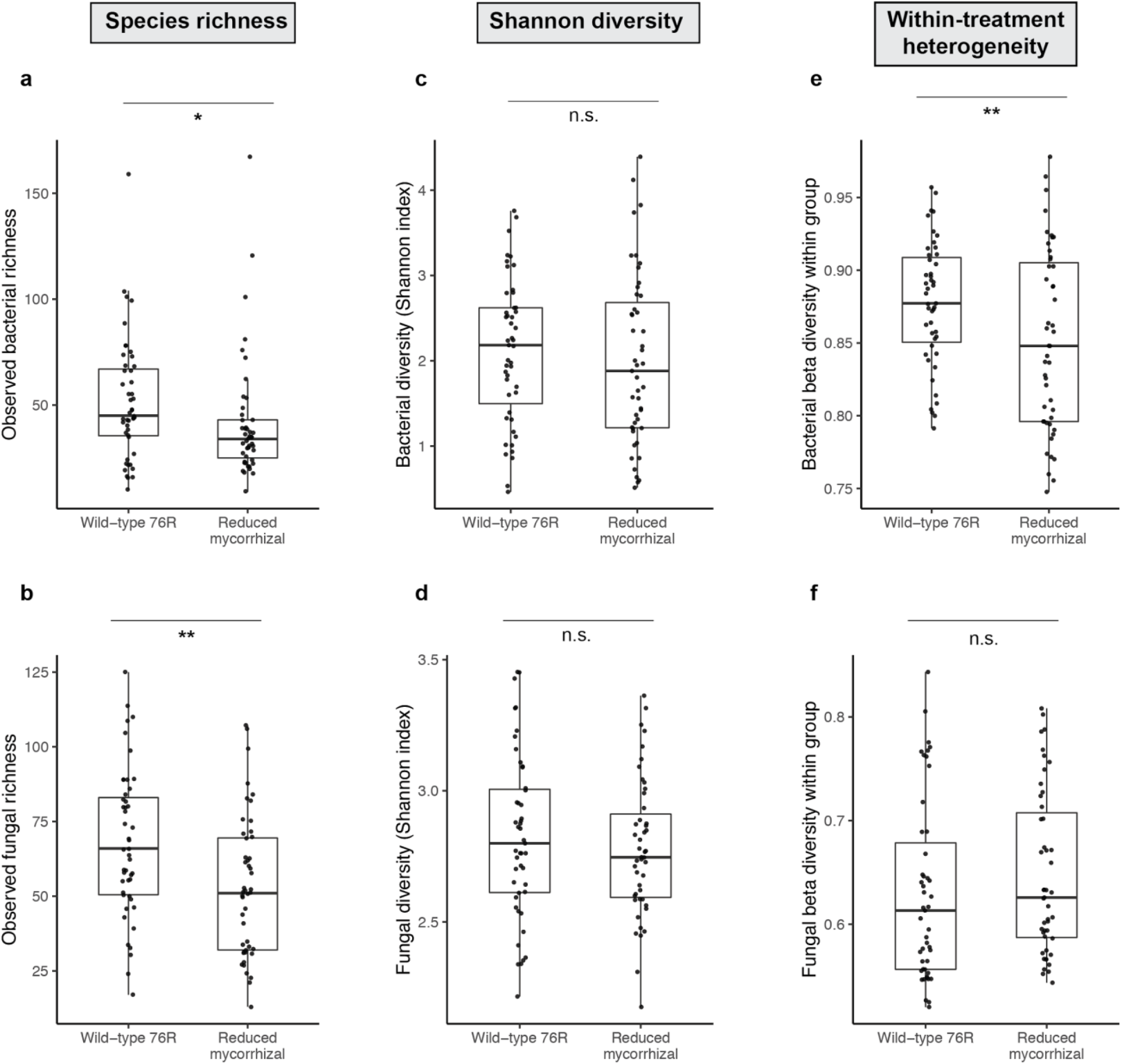
The *rmc* genotype harbors lower bacterial alpha and beta diversity, and lower fungal alpha diversity, than its wild-type counterpart. **(a**,**b)** ASV counts after removing plant sequences and contaminant sequences. Bacterial richness values were log-transformed prior to ANOVA to meet normality and homoscedasticity assumptions. **(c**,**d)** Shannon-Weaver index of combined richness and evenness. **(e**,**f)** Within-group dispersion was quantified by generating a Bray-Curtis dissimilarity matrix. Points represent the mean Bray-Curtis distance of each sample to other samples within the same plant genotype (76R or rmc).

Having observed that water limitation and disruption of mycorrhizal associations induced similar changes in bacterial community metrics (i.e. reduction in alpha and beta diversity), we next asked whether similar taxa were affected in both treatments. A permutational ANOVA on Bray-Curtis dissimilarity separated the fully watered wild-type plants from plants in deficit and/or reduced mycorrhizal conditions, while the remaining combinations of water regime and mycorrhizal associations (wild-type/deficit, *rmc*/full water, *rmc*/deficit) were not statistically different from one another (**Figure 3A-B**). To identify specific taxa affected by each treatment, we calculated Dufrêne and Legendre’s indicator value for each bacterial ASV. There was a high degree of overlap in indicator taxa for the full water and wild-type treatments, and in indicator taxa for the water-stressed and reduced mycorrhizae treatments (**Figure 3C**). We validated this result using negative binomial modeling, another method for identifying differentially abundant taxa in microbiomes, and again found substantial parallels between the effects of water regime and mycorrhizal associations on bacterial community composition (**Figure S7**). Although the water regime and mycorrhizae treatments did not induce parallel shifts in fungal community diversity metrics, we again observed substantial overlap in the distribution of individual indicator taxa (**Figure S8**).

**Figure 3.**
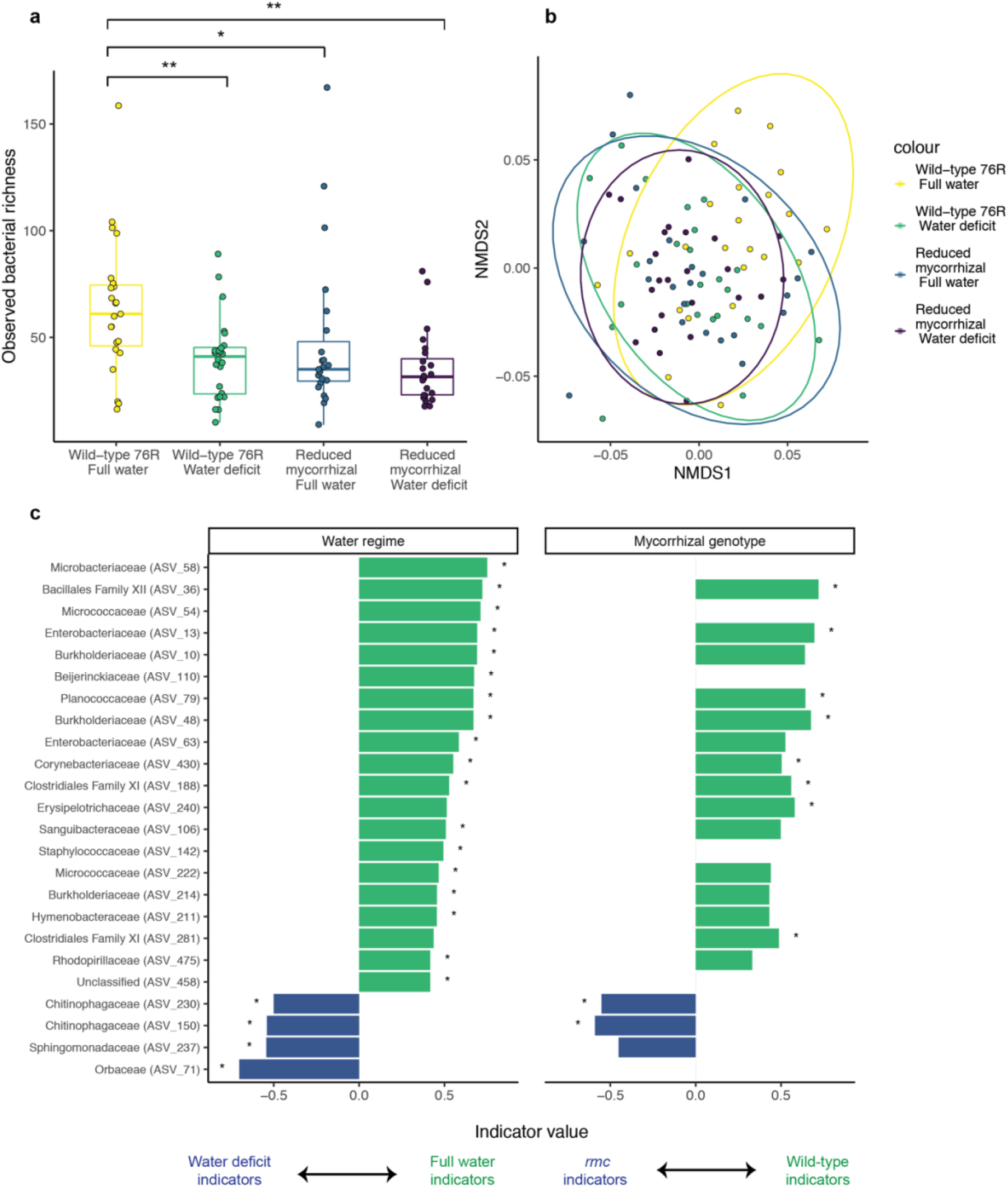
Water stress and *rmc* genotype induce parallel shifts in bacterial composition. **(a)** ASV counts after removing plant sequences and contaminant sequences. Bacterial richness values were log-transformed prior to ANOVA. **(b)** Non-metric multidimensional scaling (NMDS) ordination shows that water-stressed and *rmc* plants cluster together, away from fully watered wild-type plants. Ellipses indicate 95% confidence around clustering. **(c)** Indicator ASVs identified for water (on wild-type background) or mycorrhizae (on a full water background). Names indicate family-level classification, with ASV name in parenthesis. Values represent the square root of Dufrêne and Legendre’s indicator value. For ease of interpretation, the indicator values of opposing treatments (water deficit vs. full water, reduced mycorrhizal vs. wild-type) are displayed with opposing signs. Asterisks indicate nominal significance (p<0.05). Missing bars indicate taxa that were too prevalent in both treatments to statistically test association.

In contrast, the phosphorus fertilizer gradient did not impact bacterial alpha or beta diversity. However, increasing fertilizer concentrations were associated with higher fungal alpha diversity (F=4.802, *p*=0.032) and increased heterogeneity in fungal communities among plants (F=6.293, *p*=0.0028) (**Figure S9**).

### Cross-kingdom associations

Bacterial richness and fungal richness were highly correlated across plants (*r*=0.482, *p*<0.0001, **Figure 4A**). The relationship remained significant in an ANOVA model that controlled for the field trial treatments (water regime, mycorrhizal associations and soil fertilizer concentration) and technical batches (plant sonication batch and DNA extraction batch). Partial R^2^ calculation indicated that bacterial richness accounted for 22.4% of the variation in fungal richness in this model. Bray-Curtis dissimilarity measures based on bacterial community composition and fungal community composition were correlated as well, including when controlling for field trial treatments and technical batches (Partial Mantel statistic = 0.1563, *p*<0.0001).

**Figure 4.**
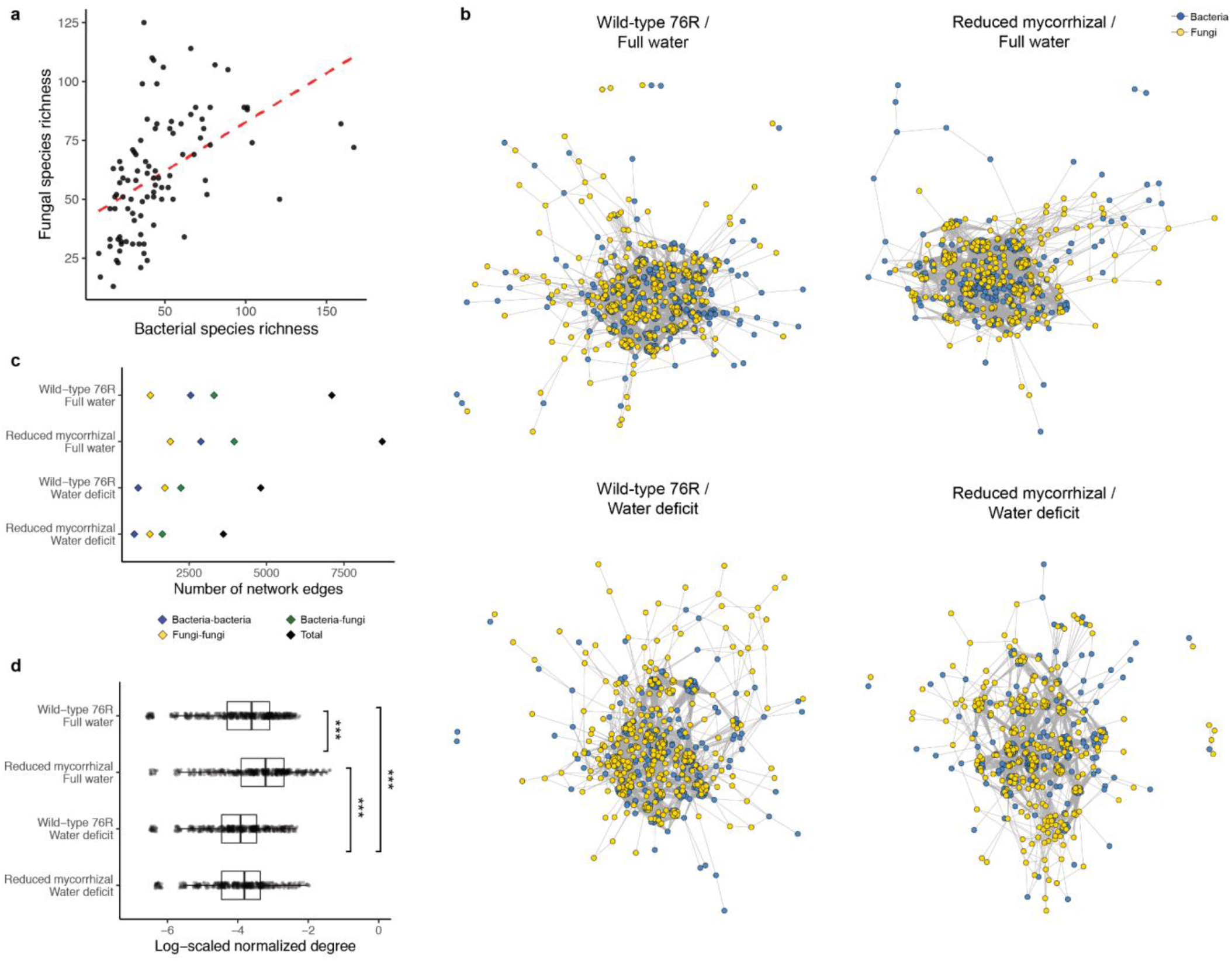
Correlations between bacterial and fungal composition. **(a)** Observed bacterial and fungal richness in the field trial, as measured by ASV counts after removing plant sequences and contaminant sequences. **(b)** Co-occurrence networks of top 500 bacterial taxa and top 500 fungal taxa, structured according to Fruchterman and Reingold’s force-directed layout algorithm. **(c)** Number of network edges (positive and negative), displayed in total and according to the taxonomic identities of the nodes. **(d)** Distribution of degree per node (number of positive and negative connections), normalized to total network size and log-transformed.

To assess the contributions of cross-kingdom associations to overall occupancy patterns, we constructed a co-occurrence network for each combination of water regime and plant genotype (**Figure 4B**). Across all treatment combinations, cross-kingdom pairs made up a larger proportion of the significant co-associations than either bacteria-bacteria or fungi-fungi pairs. Across mycorrhizal treatments, water stress decreased the average connectedness of nodes. These differences appeared to be driven by decreases in the number of bacteria-bacteria and bacteria-fungi associations in water-stressed plants (**Figure 4C-D**).

### Growth chamber experiments

We next sought to explore phyllosphere microbiome assembly and function in a more controlled setting. Microbial cells were suspended in a sterile buffer and sprayed onto three-week-old tomato plants. In the mycorrhizae experiment, wild-type and *rmc* plants in the growth chamber were sprayed with inocula from wild-type or *rmc* plants in the field, or a sterile buffer, for a fully reciprocal design. In the phosphorus experiment, plants treated with fertilizer matching the high and low extremes of the fertilizer gradient were sprayed with inocula from the corresponding positions in the field, or a sterile buffer. In both experiments, each individual sample (corresponding to one plant in the field) was divided in half and used to generate inocula for two plants, one in each growth chamber treatment.

The purpose of these experiments was twofold. First, they served as an independent and more controlled test of the treatment effects in the field trial. Pots were completely randomized with respect to treatment, and the paired design allowed direct comparison of plants that received the same inoculum but were grown in different conditions. Second, after allowing communities to establish for one week, we inoculated plants with the bacterial pathogen *Pseudomonas syringae* to test whether stressful conditions in the field had selected for microbiota that improve plant stress tolerance.

Plants in the growth chamber harbored less diverse bacterial communities than their inocula (mycorrhizae experiment: F=52.369, *p*<0.0001; phosphorus experiment: F=193.307, *p*<0.0001) and underwent a strong shift in bacterial community composition (*p*<0.0001). In the mycorrhizae experiment, bacterial communities remained more similar to their inocula when sprayed onto a plant of the same genotype than a plant of the opposite genotype (*p*=0.041). Similarly, bacterial communities in the phosphorus experiment remained more similar when the fertilizer treatment of the field plant matched that of the growth chamber plant (*p*=0.0017) **(Figure 5)**. The loss of fungal diversity in the growth chamber was much greater, and many plants were dominated by fungal ASVs that were not present in the inoculum (**Figure S10)**, suggesting that the microbiome transplant methods we used were not optimized for fungi. As such, we focused on the bacterial communities in all analyses.

**Figure 5:**
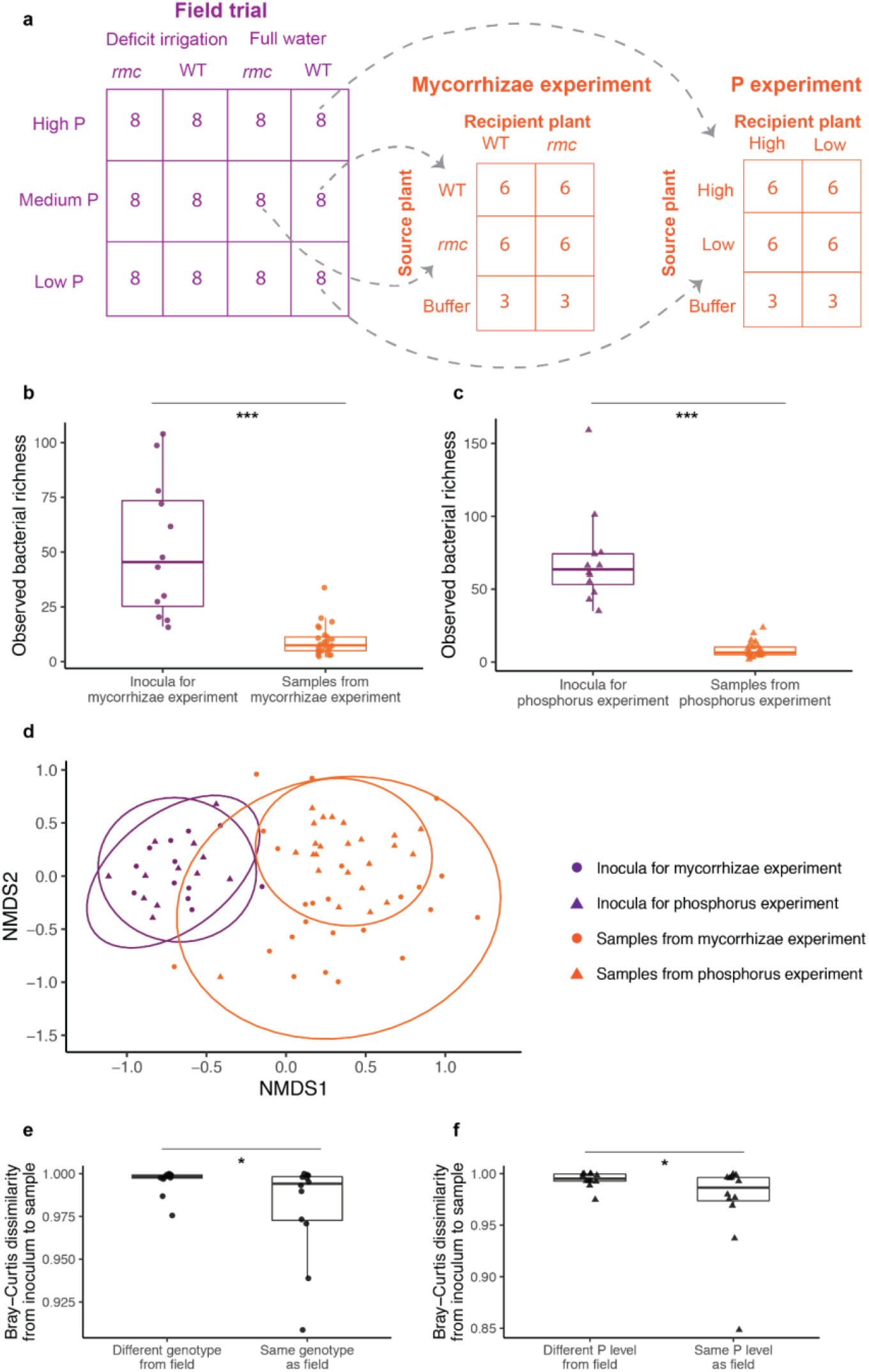
Diversity and compositional changes from field to growth chamber. **(a)** Schematic indicating communities from the field that were transplanted onto growth chamber plants. **(b**,**c)** Observed bacterial richness, as measured by ASV counts after removing plant sequences and contaminant sequences. Richness values were log-transformed prior to ANOVA. **(d)** NMDS ordination of inocula and samples from growth chamber experiments. Ellipses indicate 95% confidence around clustering. **(e**,**f)** Bray-Curtis dissimilarity index between each field plant used to generate inocula and its corresponding inoculated plants in the growth chamber. Categories indicate whether the plant in the growth chamber received the same or different experiment treatment as its source plant in the field. Each individual field plant was used to generate inocula for both growth chamber treatments, thus allowing a paired test.

The results of the growth chamber experiments generally recapitulated the effects observed in the field. In the genotype experiment, *rmc* plants harbored lower bacterial alpha and beta diversity. In the phosphorus experiment, as in the field, bacterial alpha and beta diversity were similar across treatments (**Figure S11**). Contrary to our expectations, plants inoculated with microbiota from source plants grown under matching conditions in the field did not have higher disease resistance, though wild-type plants were more susceptible to *Pseudomonas syringae* than *rmc* plants in general (**Figure S12**).

## DISCUSSION

All plants have evolved in a microbial world, yet we are only beginning to understand how microbial communities reciprocally influence and are influenced by their host. Systemic changes in nutrient and biomass allocation, defense regulation, and structural characteristics of leaves in water- and nutrient-limited plants (Zhou et al. 2019; Moles et al. 2018; Bowles et al. 2016) all suggest that soil conditions are likely to impact aboveground microbiota, with potential consequences for plant health. Here, we tested the effects of irrigation, mycorrhizae, and soil fertility on phyllosphere bacterial and fungal communities.

In the field and growth chamber, water stress and mycorrhizal disruption reduced bacterial richness and homogenized communities across host plants. Together, reduced species richness and homogenization reflect a disproportionate loss of bacterial species with low occupancy in the metacommunity (i.e. those that differentiate individual plants from one another). The phyllosphere is an environment with harsh temperature fluctuations and low relative humidity compared to the soil or rhizosphere (Knief et al. 2012; Vorholt 2012). Bacterial taxa that only colonize a small proportion of plants may be less well-adapted to these conditions, and therefore more strongly affected by stressors. Alternatively, low metacommunity occupancy tends to correlate with small local population sizes (**Figure S13**, (Gaston 1994)), which are often more vulnerable to ecological stochasticity (Shoemaker et al. 2020) or genetic drift (Lynch, Conery, and Burger 1995). In contrast, fungal community evenness increased under water stress and mycorrhizal disruption, reflecting a relative loss of dominant taxa. One explanation is that a small number of dominant taxa may comprise the majority of actively growing cells, which are more strongly affected by stressors, while rare sequences originate from dormant spores. In support of this explanation, RNA- and stable isotope-based profiling show that actively growing fungi can differ substantially from the total fungal community (Cardoso et al. 2017; Che et al. 2018) and are more sensitive to disturbance (Che et al. 2019).

In general, water stress affected bacterial communities more strongly than fungal communities, a pattern consistent with a large body of work comparing bacterial and fungal drought responses in soil (Barnard, Osborne, and Firestone 2013; de Vries et al. 2018; Ochoa-Hueso et al. 2018; Sun et al. 2020; Preece et al. 2019; Yuste et al. 2011). Interactions between bacteria and fungi may be an important part of the picture as well. We found strong positive correlations between bacterial and fungal composition that persisted when accounting for measured environmental variation. The connectedness of the multi-kingdom network decreased under water-limited conditions despite similar network sizes, largely due to decreases in the numbers of bacteria-fungi and bacteria-bacteria associations. Our ability to draw direct inferences about cross-kingdom interactions in this study is limited, as co-associations can be confounded by environmental heterogeneity or differences between measured and biologically relevant spatial scales (Blanchet, Cazelles, and Gravel 2020). However, other work has compiled many fascinating examples of interactions between bacteria and fungi (Scherlach, Graupner, and Hertweck 2013), and it will be important to see whether and how environmental stressors disrupt these interactions.

We observed a striking parallelism in the responses of bacterial taxa to water stress and the *rmc* genotype in our study, and moderate parallelism in the responses of fungi. Based on these observations, we propose that reduced soil moisture and disruption of mycorrhizae both induce systemic stress responses that alter aboveground plant physiology and microbiome composition. Both water limitation and mycorrhizal disruption can induce stomatal closure, reduce specific leaf area, and alter leaf nutrient content (Easlon and Richards 2009; Augé, Toler, and Saxton 2015; Bowles et al. 2016), altering physical and chemical characteristics of the phyllosphere habitat. Interestingly, the effect of water stress in this study was strongest in wild-type plants, and the effect of mycorrhizal disruption was strongest in well-watered plants. In fact, the combined effects of water stress and mycorrhizal disruption were not associated with additional loss in microbial biodiversity compared to either individually. This observation contrasts with previously observed interactions between irrigation and mycorrhizae, where mycorrhizal plants are typically more drought-tolerant (Al-Karaki, McMichael, and Zak 2004; Augé, Toler, and Saxton 2015). It is thus possible that mycorrhizae allow plants to access compensatory pathways that buffer their own fitness under water limitation, but affect phyllosphere microbial communities more strongly. For example, mycorrhizal tomato plants closed their stomata more quickly than non-mycorrhizal plants under drought conditions in a prior study (Lazcano, Barrios-Masias, and Jackson 2014). Rapid stomatal closure in mycorrhizal plants appeared to minimize plant biomass loss compared to non-mycorrhizal plants, but may have limited microbial access to stomata, an important source of water in the phyllosphere (Beattie and Lindow 1999; Remus-Emsermann et al. 2014). It is also possible that an interaction between water stress and mycorrhizae would have emerged if plants were sampled later in the growing season, when water stress was more severe.

Several alternative mechanisms may contribute to the effects of the belowground treatments in this study on phyllosphere microbiome composition. First, microbiome variation between wild-type and reduced mycorrhizal plants could simply reflect host genetic variation, independent of the presence of mycorrhizae. In addition to *CYCLOPS/IPD3*, which confers the reduced mycorrhizal phenotype, the *rmc* deletion spans parts of four other genes (Larkan et al. 2013). At least one of the remaining genes regulates resistance to the root pathogen *Fusarium oxysporum* (Prihatna, Barbetti, and Barker 2018), but we are not aware of any off-target effects on phyllosphere bacteria or fungi. Further, reciprocal grafting experiments revealed that the *rmc* phenotype is only generated through signals from the roots, not the leaves, of *rmc* plants (Larkan et al. 2013), and *CYCLOPS/IPD3* is not highly expressed in the leaves of other plant species (*Medicago truncatula*, (Messinese et al. 2007); *Lotus japonicus* (Yano et al. 2008)).

Another explanation for the effect of the reduced mycorrhizal genotype is that mycorrhizal colonization modifies host immune signaling both locally and systemically (J. Liu et al. 2007), tending to increase resistance to necrotrophic pathogens and decrease resistance to biotrophic pathogens (Azcón-Aguilar et al. 2009). This may explain why wild-type plants were more susceptible to *Pseudomonas syringae* in the growth chamber, but it is unclear whether commensal microbiota on the leaf surface interact closely enough with plant cells to be regulated by immune signaling. Bacteria can often grow to high densities on the leaf surface with minimal changes to plant gene expression (Vogel et al. 2016), and the results of studies that directly tested for immune control of phyllosphere microbiome composition have been mixed (Chen et al. 2020; Bodenhausen et al. 2014).

Soil is thought to be one of many sources of dispersal to the phyllosphere (Copeland et al. 2015), so changes in the soil community could affect the phyllosphere community, independent of systemic plant modifications. However, the taxonomic patterns of enrichment and depletion that we observed in the phyllosphere were markedly different from those previously observed in soil and rhizosphere studies. In soil and rhizosphere habitats, drought is commonly associated with a relative enrichment of *Actinobacteria*, and depletion of *Acidobacteria, Proteobacteria*, and *Bacteroidetes* (Naylor et al. 2017; Barnard, Osborne, and Firestone 2013; Xu et al. 2018). In contrast, we did not observe significant enrichment in any families of Actinobacteria (though several were depleted under water-limited conditions), while several *Proteobacteria* and *Bacteroidetes* families were enriched under water stress (See **Table S4** for annotated list of enriched and depleted taxa). Additionally, we would not expect to see parallel responses to water stress and the *rmc* genotype if the effect were driven by dispersal. While mycorrhizae can alter surrounding soil communities, their effects tend to be relatively minor and/or highly localized (Marschner and Baumann 2003; Cavagnaro et al. 2006).

Although water regime and mycorrhizal associations were the strongest predictors of phyllosphere microbiome composition in our study, we observed some effects of fertilization. Fungal diversity and heterogeneity increased along the fertilization gradient, and bacterial communities remained more similar from the field to the growth chamber if they were transplanted onto a plant grown under the same nutrient conditions. Interestingly, these effects manifested aboveground even though fertilization did not appear to alter phosphate availability in the soil. It is possible that even though net mineralization of the organic P fertilizer was not detected, P cycling and availability was sufficiently altered as to affect plant physiology or soil composition in ways that subsequently impacted aboveground microbial communities.

In contrast to previous work in the rhizosphere microbiome (Carrión et al. 2019; H. Liu et al. 2021), transplanting microbial communities from stressful conditions onto plants in the growth chamber did not improve disease resistance, so we cannot conclude whether stress-induced compositional shifts in phyllosphere microbiota are adaptive for the host plant. While it is always possible that other, unmeasured traits were improved by microbiome transplant, our results may reflect growing evidence that host selection is weaker in the phyllosphere than the rhizosphere (Wagner et al. 2020).

Our study demonstrates the effects of soil conditions on the diversity, composition, and distribution of aboveground microbiota. We observed strikingly parallel effects of water stress and mycorrhizal disruption despite differences in the nature of these perturbations (biotic or abiotic), yet these changes were distinct from the enrichment patterns previously observed in the soil and rhizosphere. Taken together, our findings are unlikely to be explained by off-target effects of the *rmc* mutation or dispersal from soil. Rather, they indicate that systemic stress responses in host plants alter selection on aboveground microbial communities. Our study highlights the importance of soil conditions for aboveground microbiome community assembly and biodiversity.

## Acknowledgments

We would like to thank the University of California, Davis Student Farm for access to the fields, members of the Koskella and Bowles labs for helpful discussion and comments, and Microbiome Insights, Inc. for their role in sequencing efforts. Support for this work was provided by the Army Research Office (W911NF-17-1-0231 to AG), the National Science Foundation (NSF Graduate Research Fellowships to RD, CH, AG), the University of California, Berkeley (Berkeley Fellowships to YS, CH), the Society for the Study of Evolution (Grant 047408 to RD), the Hellman Fellows Fund (Hellman Fellows Award to BK) and the Marian E. Koshland Integrated Natural Sciences Center (KINSC Summer Scholarship to GK).

