## Supplementary Information for "Water Stress and Disruption of Mycorrhizae Induce Parallel Shifts in Phyllosphere Microbiome Composition"

**Supplemental Material**

*Field trial preparation and sampling*

Field preparation included spading and bed formation (1.5 m wide from furrow to furrow), followed by feather meal (12 − 0 − 0) banded in at a rate of 40 kg N ha− 1. Experimental plots were established in eight experimental beds with rows of buffer plants between each experimental bed.

Transplants were grown in the Westside Transplant facility in Winters, CA. After 9 wk. under certified organic management, seedlings were transplanted on the bed center by hand on 28 April 2019, followed by 1.9 cm of water applied via a single surface drip line in the center of each bed. Transplants that died due to a fungal blight (~100 plants) were replaced in the first three weeks of the experiment.

Soil samples were taken at three depths (0-15cm, 15-30cm, 30-60cm) at transplant, mid-season (52 days after transplant) and harvest (116 days after transplant).

*Growth chamber experiments*

Seeds were sterilized by swirling in 2.7% bleach for 20 minutes, then washed 3 times with sterile water. 4 seeds per plate were germinated on 1% water agar. Plates were wrapped in aluminum foil for 7 days and stored at 21°C. After 1 week, plants were transferred to individual pots containing Sunshine #4 potting mix, where they were maintained at a 15 h day:9 h night light cycle.

When plants were three weeks old, field communities were used to generate inocula. Each sample from the field was divided in half, then diluted to $10^4$ CFU/mL in 10 mM $MgCl_2$. The surfactant Silwet-L-77 was added at a concentration of 0.1 µL autoclaved Silwet per 1 mL inocula to facilitate microbial adhesion onto the leaf surface. Each plant was sprayed with 4 mL of inoculum using a 15 mL conical tube fitted with a spray cap. Each plant was sprayed from all angles inside an autoclave bag to contain microbes.

One week after spraying, a frozen stock of *Pseudomonas syringae* pathovar tomato PT23 was grown overnight in King's Broth at 28°C for experimental infections. The overnight culture was diluted in 10 mM $MgCl_2$ to an optical density ($OD_{600}$) of 0.0002. At 24 hours post-infection, 3 hole punches (6-mm diameter) were taken from each inoculated leaf, for a total of 9 leaf discs per plant. Leaf discs were placed in a FastPrep tube with 1 uL 10 mM $MgCl_2$ and two sterile ceramic beads, then homogenized in the FastPrep-24 5G sample disruption instrument at 4.0 m/s for 40 seconds.

The rest of the aboveground plant tissue was placed in a 15 mL tube. The tube was filled with 10 mM $MgCl_2$ and submerged in a Branson M5800 sonicating water bath for 10 minutes to gently dislodge microbial cells from the leaf surface. Leaf wash was centrifuged at 3500 x *g* for 10 minutes, resuspended in 600 uL 10 mM $MgCl_2$, and frozen immediately at –20°C for sequencing.

*Droplet digital PCR reaction conditions and primers*

*Pseudomonas* abundance was quantified using the Bio-Rad QX200™ Droplet Digital PCR system. Reaction mixtures were prepared with 11 µL Bio-Rad ddPCR Supermix for Probes (no dUTP), 1.1 µL probe (see below), 4.9 µL molecular grade water, and 5 µL homogenized leaf material. Samples were randomized on the ddPCR plate, with a negative control (replacing the

5 µL template with molecular grade water) in the last well of each column. Droplets were generated according to manufacturer instructions, and the following PCR conditions were used for amplification: 95°C for 10 minutes; 40 cycles of the following: (94°C for 30 seconds, 60°C for 1 minute, 72°C for 1 minute); 98°C for 10 minutes; hold at 12°C. Positive droplet thresholds were set by column, with fluorescence values in the range of the negative control classified as negative. ddPCR primers were adapted from *Pseudomonas* qPCR assays and target a hypervariable region of the 16S rRNA gene (Bergmark et al. 2012): Forward 5'-ACTTTAAGTTGGGAGGAAGGG-3'; reverse 5'-ACACAGGAAATTCCACCACCC-3', probe TGCCAGCAGCCGCGG.

Total bacterial abundance was quantified using the Bio-Rad QX200™ Droplet Digital PCR system. Reaction mixtures consisted of 11 µL EvaGreen Supermix, 0.22 µL forward primer (see below), 0.22 µL reverse primer, 5.56 µL molecular grade water, and 5 µL leaf wash. Samples were randomized on the ddPCR plate, with a negative control in the last well of each column. Droplets were generated according to manufacturer instructions, and the following PCR conditions were used for amplification: 95°C for 10 minutes; 40 cycles of the following: (95°C for 30 seconds, 55°C for 30 seconds, 72°C for 2 minutes), 4°C for 5 minutes, 90°C for 5 minutes, hold at 12°C. Positive droplet thresholds were set by column, with fluorescence values in the range of the negative control classified as negative. Conserved sequences in the V3-V4 regions of the 16S rRNA gene were targeted using the following primers: Forward 5'-CCTACGGGNBGCASCAG-3'; reverse 5'-GACTACNVGGGTATCTAATCC-3'.

*Amplicon sequencing workflow and analysis*

The V4 region of the 16S rRNA gene was amplified using 515F/806R primers, with peptide nucleic acids (PNAs) added to limit amplification of plant chloroplast and mitochondrial DNA (Lundberg et al. 2013). The eukaryotic ITS2 gene was amplified using the primers (5'-CCTCCGCTTATTGATATGC-3', 5'-CCGTGARTCATCGAATCTTTG-3'). Reads were analyzed using the recommended DADA2 pipeline with the following parameters. Reads with Ns or >2 "expected errors" (calculated by DADA2 based on quality scores) were removed. 16S reads were truncated at 240 base pairs in the forward direction and 140 base pairs in the reverse direction, or at the first position with a quality score <2. ITS reads shorter than 50 base pairs were removed, or were truncated at the first position with a quality score <2. Amplicon sequence variants (Callahan, McMurdie, and Holmes 2017) were inferred after training the core denoising algorithm on error rates. DADA2 was then used to merge paired reads, on the condition that forward and reverse reads overlap by at least 12 bases. Chimeric sequences were identified as exact reconstructions of segments of two more abundant "parent" sequences, and removed from further analysis. After filtering plant-derived sequences and sequences prevalent in negative controls, the dataset consisted of 1579 bacterial amplicon sequence variants (ASVs) and 1397 fungal ASVs.

**Table S1. Fertilizer gradient in field trial.** Bold values indicate sampling scheme.

| **Position in gradient** | **Fertilizer added ($g/m^2$)** | **Fertilizer added ($g/m^3$)** (assuming penetration depth of 20 cm) |
|---|---|---|
| **1, 2** | **0** | **0** |
| 3 | 40 | 200 |
| 4 | 80 | 400 |
| 5 | 120 | 600 |
| 6 | 160 | 800 |
| 7 | 200 | 1000 |
| **8** | **400** | **2000** |
| **9** | **600** | **3000** |
| 10 | 800 | 4000 |
| **11, 12** | **1000** | **5000** |

**Table S2. Full results of ANOVA models of bacterial and fungal richness.**

| **Bacterial ASV richness** (log-transformed) | | | | | |
|---|---|---|---|---|---|
| **Treatment** | **Df** | **Sum of squares** | **Mean square** | **F value** | **p-value** |
| **Water regime** | **1** | **2.384** | **2.384** | **10.468** | **0.0018** |
| **Genotype** | **1** | **1.185** | **1.185** | **5.203** | **0.026** |
| Fertilizer concentration | 1 | 0.387 | 0.387 | 1.697 | 0.197 |
| Plant sonication batch | 12 | 2.729 | 0.227 | 0.998 | 0.459 |
| **DNA extraction batch** | **6** | **6.765** | **1.127** | **4.951** | **0.00027** |
| **Fungal ASV richness** | | | | | |
| **Treatment** | **Df** | **Sum of squares** | **Mean square** | **F value** | **p-value** |
| Water regime | 1 | 388.5 | 388.5 | 0.902 | 0.345 |
| **Genotype** | **1** | **4919.1** | **4919.1** | **11.417** | **0.0011** |
| **Fertilizer concentration** | **1** | **2069.1** | **2069.1** | **4.802** | **0.032** |
| Plant sonication batch | 12 | 8802.5 | 733.5 | 1.702 | 0.084 |
| **DNA extraction batch** | **6** | **10317.9** | **1.19.7** | **3.991** | **0.0017** |

**Table S3. Full results of ANOVA models of bacterial and fungal Shannon diversity.**

| **Bacterial Shannon diversity index** | | | | | |
|---|---|---|---|---|---|
| **Treatment** | **Df** | **Sum of squares** | **Mean square** | **F value** | **p-value** |
| Water regime | 1 | 3.047 | 3.047 | 4.990 | 0.029 |
| Genotype | 1 | 0.325 | 0.325 | 0.533 | 0.468 |
| Fertilizer concentration | 1 | 0.076 | 0.076 | 0.124 | 0.726 |
| Plant sonication batch | 12 | 26.670 | 2.222 | 3.640 | 0.00028 |
| DNA extraction batch | 6 | 4.760 | 0.793 | 1.299 | 0.268 |
| **Fungal Shannon diversity index** | | | | | |
| **Treatment** | **Df** | **Sum of squares** | **Mean square** | **F value** | **p-value** |
| **Water regime** | **1** | **0.5262** | **0.5262** | **7.966** | **0.0062** |
| Genotype | 1 | 0.046 | 0.046 | 0.690 | 0.409 |
| Fertilizer concentration | 1 | 0.040 | 0.040 | 0.610 | 0.437 |
| Plant sonication batch | 12 | 1.336 | 0.111 | 1.685 | 0.088 |
| **DNA extraction batch** | **6** | **0.973** | **0.162** | **2.453** | **0.032** |

**Table S4. Bacterial families identified in enrichment analyses.**

| Family name | Water response | Genotype response | Implicated in drought? | Implicated in disease? |
|---|---|---|---|---|
| Sphingobacteriaceae | Enriched in FW | Enriched in 76R | Promotes plant growth during drought (Rolli et al. 2015); depleted in drought in soil (Meisner et al. 2018); dominates high moisture grass seedling environments (Chen et al. 2020); Sphingobacteriaceae decreased in relative abundance in soil with a history of drought and only increased when the soil was re-wetted (Meisner et al. 2018). | |
| Burkholderiaceae | Enriched in FW | N/A | Promoted plant growth during drought in combination with AMF in soil (Tallapragada, Dikshit, and Seshagiri 2016); dominates low moisture grass seedling environments (Chen et al. 2020). | *Burkholderia andropogonis* causes bacterial leaf stripe on sorghum. After sowing infected seeds, bacteria spread through the vascular system in seedlings and developing plants, penetrating into emerging and ripening seeds. The indirect vector is the flea beetle *P.vuttula* (Kaplin 2019). |
| Caulobacteraceae | Enriched in FW | Enriched in 76R | | After white fly infestation on peppers above ground, this family is one of the most abundant in the rhizosphere (Kong et al. 2016); after subterranean white grub infestation, caulobacteraceae relative abundance increased in peanut rhizosphere (Geng et al. 2018). |
| Hymenobacteraceae | Enriched in FW | Enriched in 76R | Enriched in alpine wetland soil as opposed to alpine forest soil (Wang et al. 2020); more prevalent in high moisture grass seedling environments than low moisture seedling environments (Chen et al. 2020). | |
| Bacilli Family_XII | Enriched in FW | Enriched in 76R | *Bacillus* strains are very tolerant and able to improve plant's resistance to drought (Mathur and Roy 2021); Also, *Bacillus* strains are capable to enhance number of tillers and spikelets of wheat (Raheem et al. 2018); *Bacillus* strains confer tolerance to osmotic stress in arabidopsis seedlings (Ghosh, Gupta, and Mohapatra 2019). | *Bacillus* strains can play a key role as biocontrol by producing peptide antibiotics (Verma et al. 2016). |
| Sanguibacteraceae | Enriched in FW | Enriched in 76R | | |

| | | | | |
|---|---|---|---|---|
| Planococcaceae | Enriched in FW | Enriched in 76R | *Planococcus salinarum* BCZ23 is thermal-drought tolerant because it can grow at high temperatures and moisture stress conditions (Verma et al. 2016). | |
| Corynebacteriaceae | Enriched in FW | Enriched in 76R | Application of cell extracts of Corynebacteria as a foliar spray at the seedling stage and the panicle initiation stage significantly combat the drought-induced physiological stresses of rice and increases water content of rice in drought conditions (Bowya and Balachandar 2020). | *Corynebacterium pseudotuberculosis* is an animal intracellular pathogen (Soares et al. 2013). |
| Enterobacteriaceae | Enriched in FW | Enriched in 76R | *Enterobacter* sp. FD17, endophytic bacteria, improved maize physiological parameters under drought conditions when inoculated on plants, but not as well as *Burkholderia phytofirmans* PsJN (Naveed et al. 2014); Some Enterobacter sp. are plant growth promoting rhizobacteria (enhance the productivity of crops and protect them from abiotic stresses) and they are 1-aminocyclopropane-1-carboxylic acid (ACC) deaminase containing and might have the potential to alleviate drought stress (Danish et al. 2020); Enterobacteriaceae had absence of growth in a medium with reduced water availability in cacti rhizosphere (Kavamura et al. 2013). | Bacterial Soft Rot Disease on Dragon Fruit (*Hylocereus* spp.) Caused by *Enterobacter cloacae* in Peninsular Malaysia (Masyahit et al. 2009); Same bacteria causes disease in papaya (Nishijima 1987); *Enterobacter asburiae* BQ9 promotes tomato plant growth and induce resistance to Tomato yellow leaf curl virus (Li et al. 2016). |
| Clostridiales Family_XI | Enriched in FW | Enriched in 76R | *Clostridiales* was found as a main drought sensitive species in grapevine rhizosphere (Zhang et al. 2019). | |
| Paenibacillaceae | Enriched in FW | Enriched in 76R | More prevalent in high moisture grass seedling environments than low moisture seedling environments (Chen et al. 2020); Paenibacillaceae had absence of growth in a medium with reduced water availability in cacti rhizosphere (Kavamura et al. 2013). | |
| Erysipelotrichaceae | Enriched in FW | Enriched in 76R | Wild black howler monkey gut microbiota was characterized with the genera sequenced - *solobacterium* during the dry time when they eat fruit (Amato et al. 2015). | |
| Staphylococcaceae | Enriched | N/A | *Staphylococcus devriesei* was | Some *Staphylococcaceae* are |

| | | | | |
|---|---|---|---|---|
| | in FW | | found to be a thermo-drought tolerant bacteria because it can grow at high temperatures and moisture stressed environments (Verma et al. 2016). | closely related to and considered *Bacillus* derived genera and can play a key role in biocontrol by producing peptide antibiotics (Verma et al. 2016); gymnosperms - *Juniperus sabina* and *Cephalotaxus harringtonia* have intense antibacterial ethanol extract responses from their leaves and shoots to the Staphylococcaceae family. Methicillin resistant *Staphylococcus aureus* is one of the commonest recognized antibiotic-resistant bacteria (Zazharskyi et al. 2020). |
| Microbacteriaceae | Enriched in FW | N/A | Abundant on seeds of drought tolerant wheat lines in drought conditions and rainfed conditions (Hone et al., n.d.). | |
| Beijerinckiaceae | Enriched in FW | N/A | *Beijerinckiaceae* shows a statistically meaningful cluster shift in response to drought (Jang et al. 2020). | |
| Chitinophagaceae | Enriched in D (w 76R) OR FW (w rmc) | N/A | Chitinophagaceae increased in relative abundance in soil with a history of drought (Meisner et al. 2018); abundant on seeds of both drought susceptible and drought tolerant wheat lines in drought conditions and not in rainfed conditions (Hone et al., n.d.). | Endophytic community of *Chitinophagaceae* and *Flavobacteriaceae* stains harbor chitinase genes and biosynthetic gene clusters encoding non-ribosomal peptide synthetases and polyketide synthases that were found to suppress fungal root disease for sugar beets (Carrión et al. 2019). |
| Nocardiaceae | Enriched in D (w 76R) OR FW (w rmc) | N/A | | *Nocardia* species exhibited antibacterial activity against a tested pathogen in the rhizosphere (Adegboye and Babalola 2016). |
| Diplorickettsiaceae | Enriched in D | N/A | | *D. massiliensis* has been identified as a possible human tick-borne pathogen (Merenstein, Ward, and Allen 2020). |
| Archangiaceae | Enriched in FW | Enriched in rmc | Genera of Archangianceae (though not the one we sequenced), *Myxococcales*, were found persistently depleted in rhizosphere and endosphere of drought stressed rice plants (Santos-Medellin et al., n.d.). | |
| Neisseriaceae | Enriched in D | Enriched in rmc | | The *Neisseriaceae* family of bacteria causes a range of diseases including meningitis, septicaemia, gonorrhoea and |

| | | | | |
|---|---|---|---|---|
| | | | | endocarditis, and extracts haem from haemoglobin as an important iron source within the iron-limited environment of its human host (Wong et al. 2015). |
| Atopobiaceae | Enriched in FW | Enriched in rmc | | |
| Sphingomonadaceae | Enriched in D | Enriched in rmc | Drought treatment increased the abundance of *Sphingomonas* in the peanut rhizosphere (Dai et al. 2019). | |
| Orbaceae | Enriched in D | N/A | | Orbaceae species were enriched in the midgut of diseased honey bees from jujube flower disease (Ma et al. 2020) |

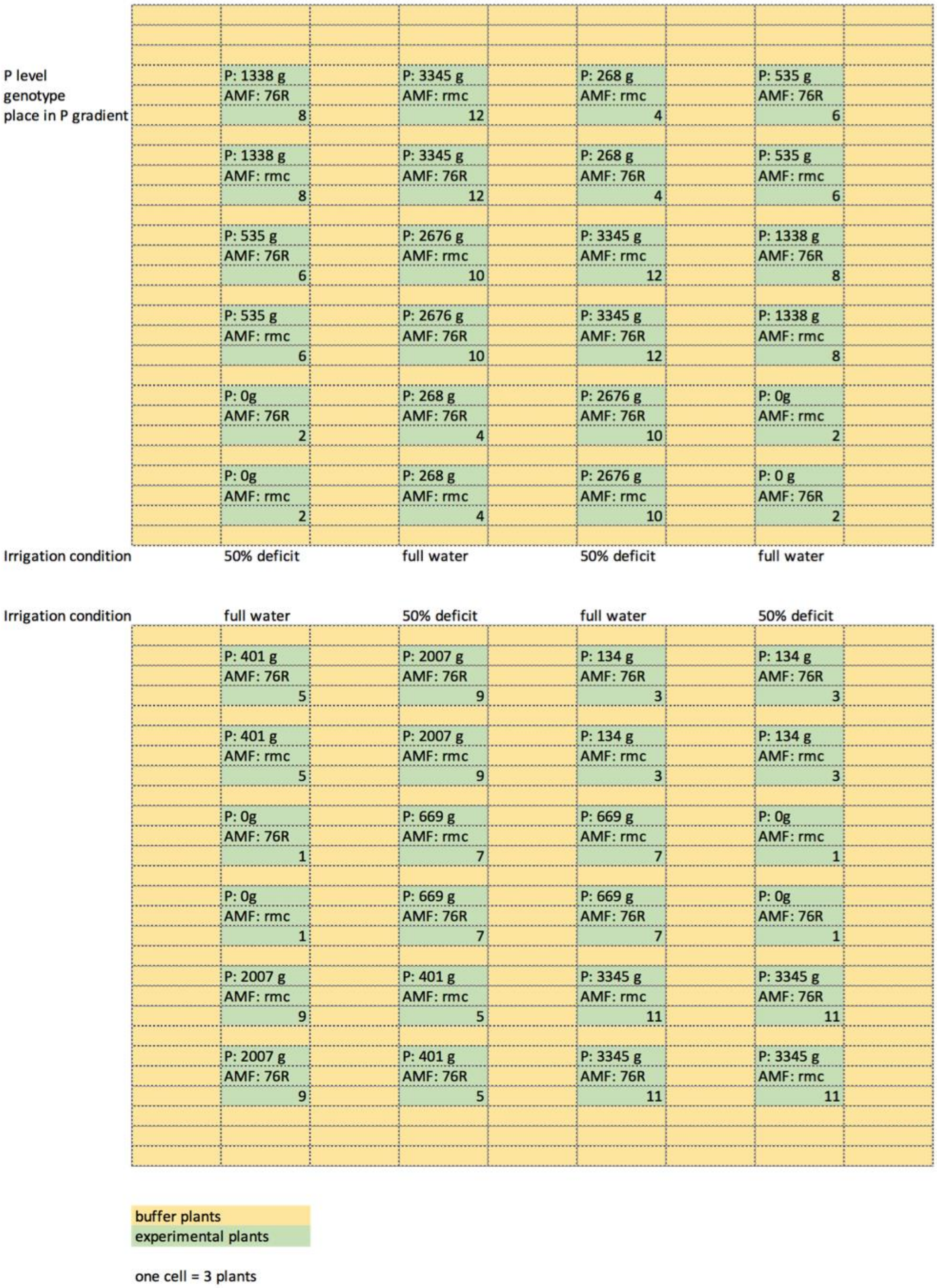

**Figure S1. Experimental design for the field trial.** Plots shown in green included 10 plants and were experimentally manipulated as described within the plot.

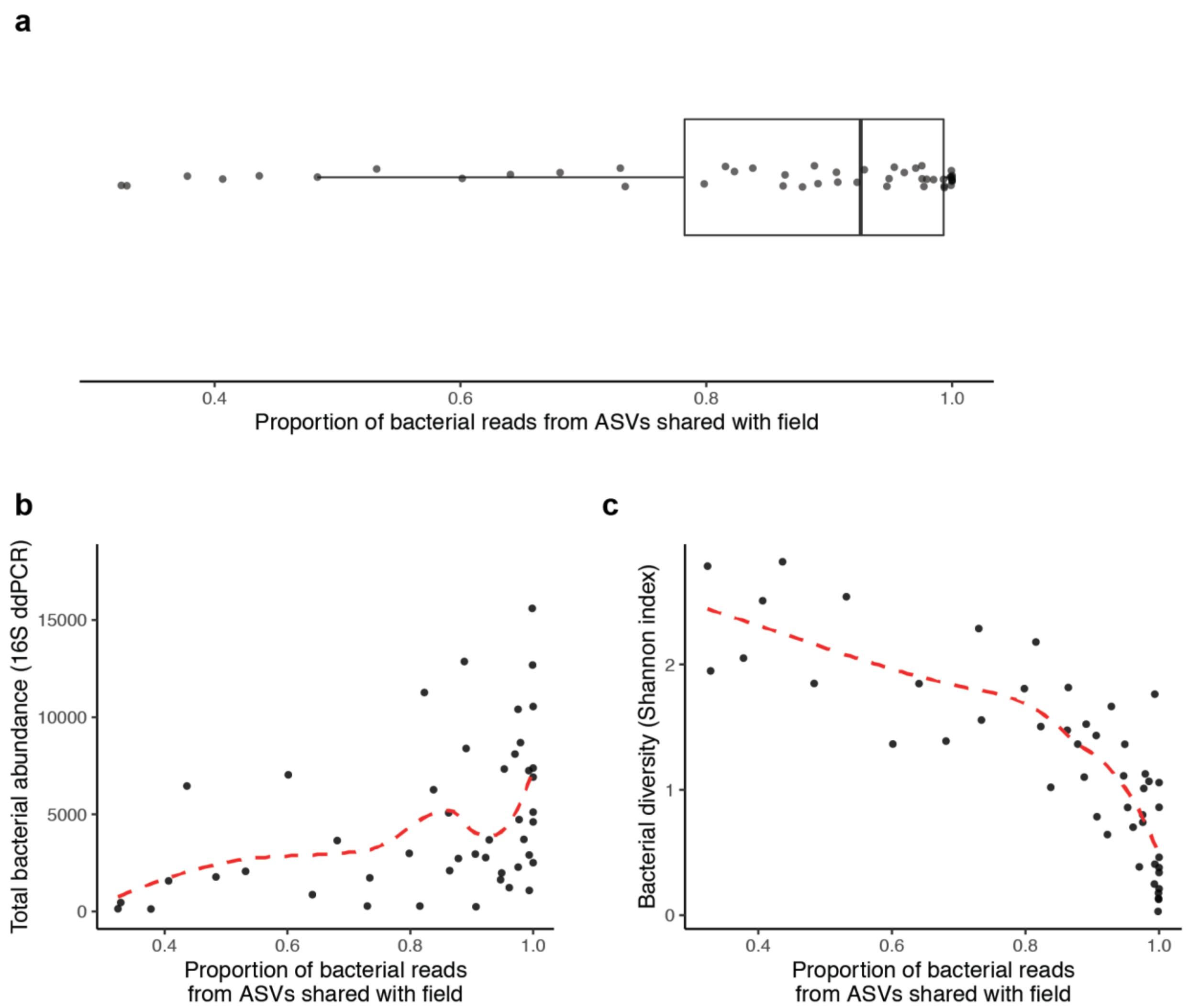

**Figure S2. Colonization of growth chamber plants by field bacterial communities. (a)** Bacterial communities on plants in the genotype and phosphorus experiments were composed primarily of taxa from the field trial (median = 92.6%), but in a subset of plants, a large proportion of reads came from taxa unique to the growth chamber. **(b)** Plants with a large proportion of growth-chamber-unique reads had low overall bacterial abundance, suggesting that the inocula failed to grow to high densities. **(c)** Plants with a large proportion of growth-chamber-unique reads had high diversity despite low bacterial abundance, suggesting that occasional contamination by growth-chamber-unique taxa confounded alpha diversity estimates.

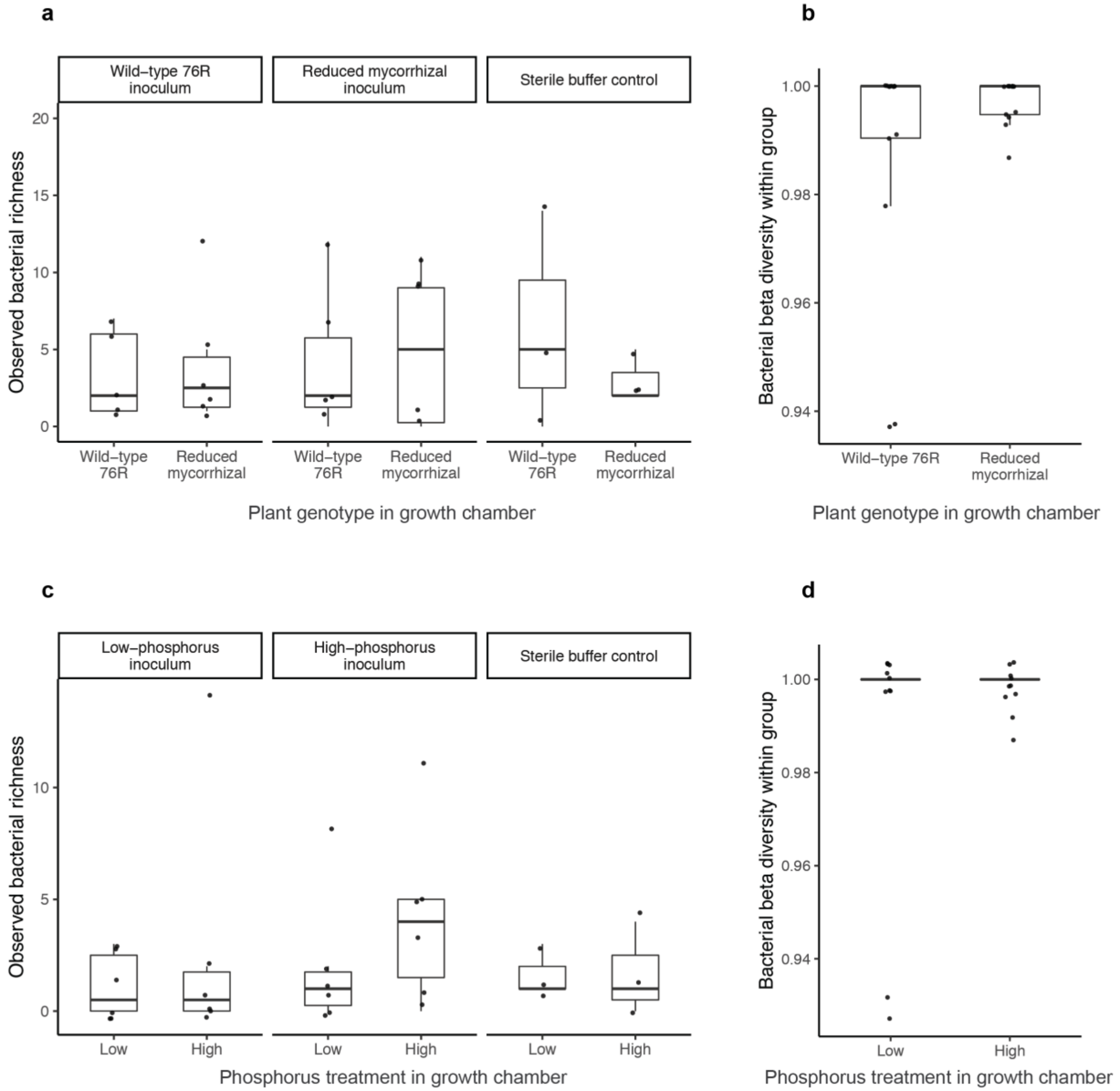

**Figure S3. Growth-chamber-unique taxa are not distributed according to experimental treatments in the growth chamber. (a)** Observed bacterial richness in genotype experiment, as measured by ASV counts of taxa unique to the growth chamber. **(b)** Within-group dispersion of bacterial communities was quantified by generating a Bray-Curtis dissimilarity matrix of taxa unique to the growth chamber. Points represent the mean Bray-Curtis distance of each sample to other samples of the same genotype (76R or reduced mycorrhizal). **(c)** Observed bacterial richness in phosphorus experiment, as measured by ASV counts of taxa unique to the growth chamber. **(d)** Within-group dispersion of bacterial communities was quantified by generating a Bray-Curtis dissimilarity matrix of taxa unique to the growth chamber. Points represent the mean Bray-Curtis distance of each sample to other samples within the same treatment group (low, high).

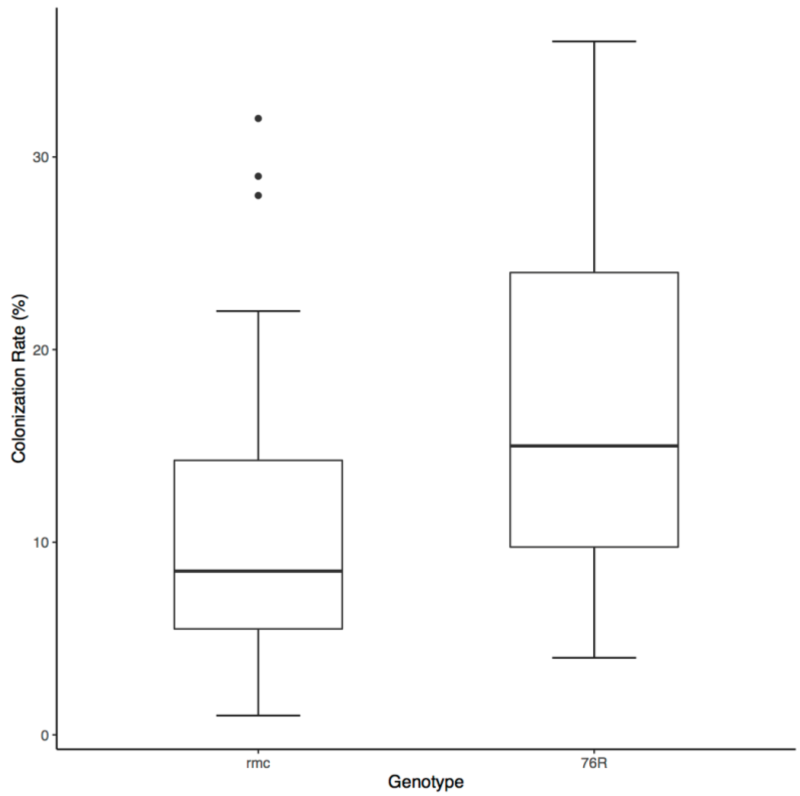

**Figure S4. Root AMF colonization rates of plants in field trials by genotype.** Wildtype plants (76R) show significantly higher root colonization by AMF than reduced mycorrhizal mutants (*rmc*).

a

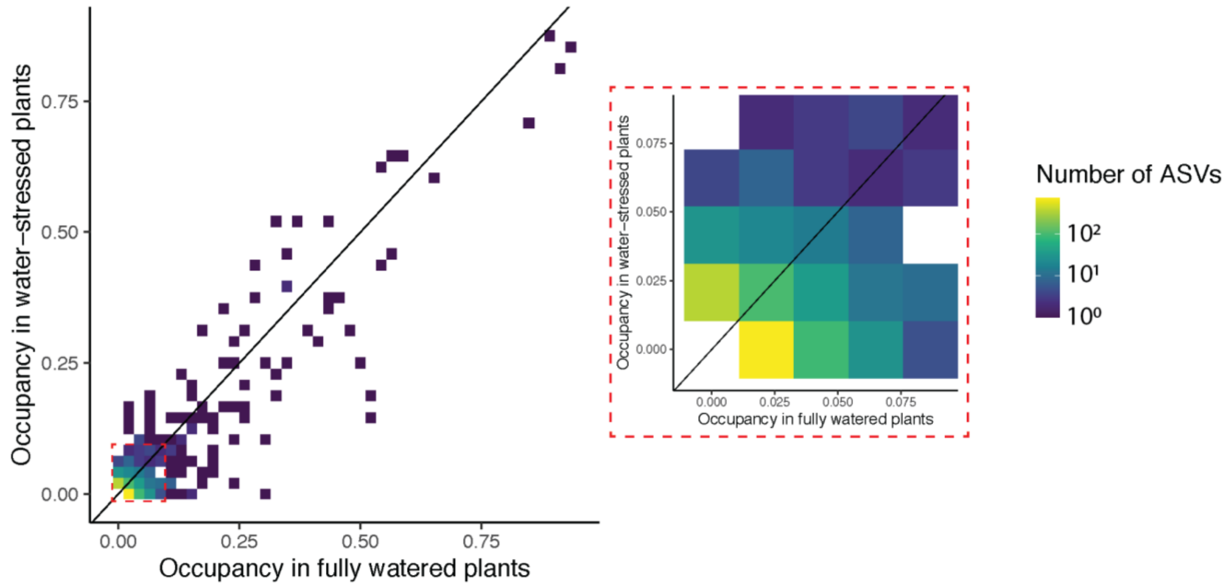

b

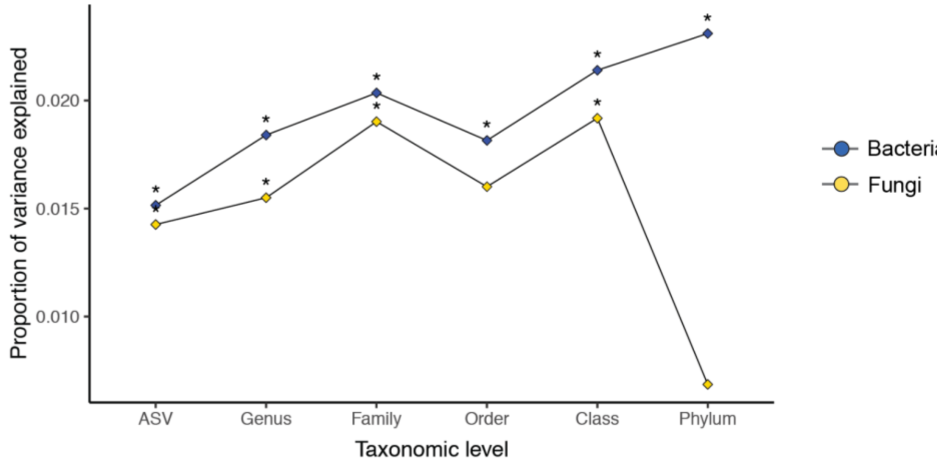

**Figure S5. Effects of water stress are strongest on rare bacterial taxa and persist at broad taxonomic levels. (a)** Tile plot describing the relationship between the proportion of fully watered plants and the proportion of drought plants occupied (relative abundance > 0) by each bacterial ASV. Inset depicts bacterial ASVs that were detected on fewer than 10% of plants in either treatment. Many of these ASVs were less frequently detected in the drought treatment than in the full water treatment (as indicated by brighter colors below the diagonal). **(b)** Proportion of variance in bacterial or fungal community composition explained by water regime in a model including plant genotype, fertilizer concentration, plant sonication batch, and DNA extraction batch as additional covariates. Asterisks indicate where drought explained a significant portion of variation in a PERMANOVA test.

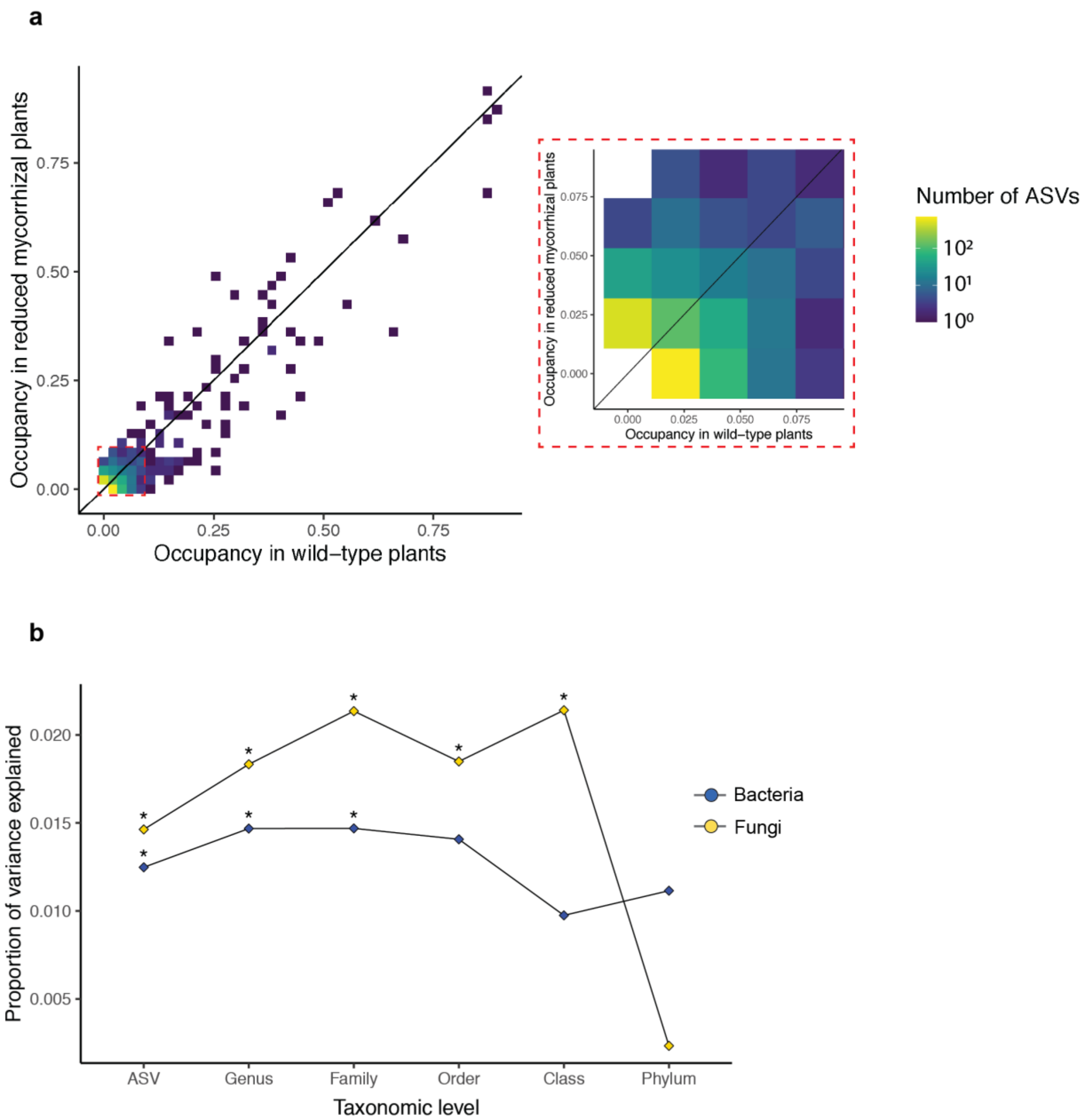

**Figure S6. Effects of the *rmc* genotype are strongest on rare bacterial taxa and persist at broad taxonomic levels. (a)** Tile plot describing the relationship between the proportion of wild-type plants and the proportion of reduced mycorrhizal plants occupied (relative abundance > 0) by each bacterial ASV. Inset depicts bacterial ASVs that were detected on fewer than 10% of plants in either treatment. Many of these ASVs were less frequently detected in reduced mycorrhizal plants than in wild-type plants (as indicated by brighter colors below the diagonal). **(b)** Proportion of variance in bacterial or fungal community composition explained by mycorrhizal genotype in a model including water regime, fertilizer concentration, plant sonication batch, and DNA extraction batch as additional covariates. Asterisks indicate where plant genotype explained a significant portion of variation in a PERMANOVA test.

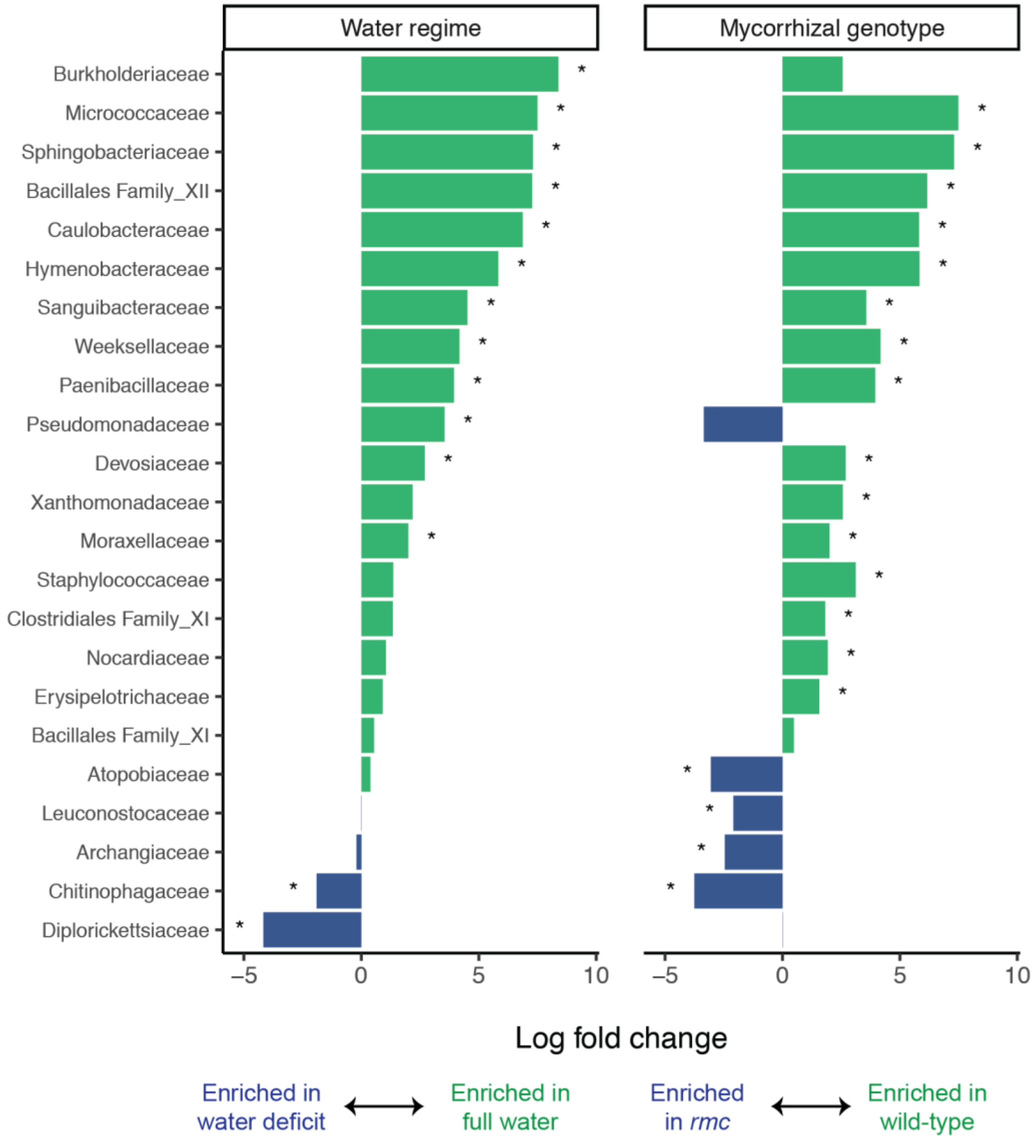

**Figure S7. Negative binomial modeling identifies parallel shifts in bacterial community composition in water stress and *rmc* genotype.** The negative binomial model implemented in the package *edgeR* was used to calculate log fold differences in abundance of bacterial families across water regimes (on a wild-type background) or plant genotypes (on a full water background). Significance was assessed for each family using an analog of Fisher's exact test adapted for overdispersed data, then corrected for multiple testing. Asterisks indicate significance after multiple testing (FDR<0.01).

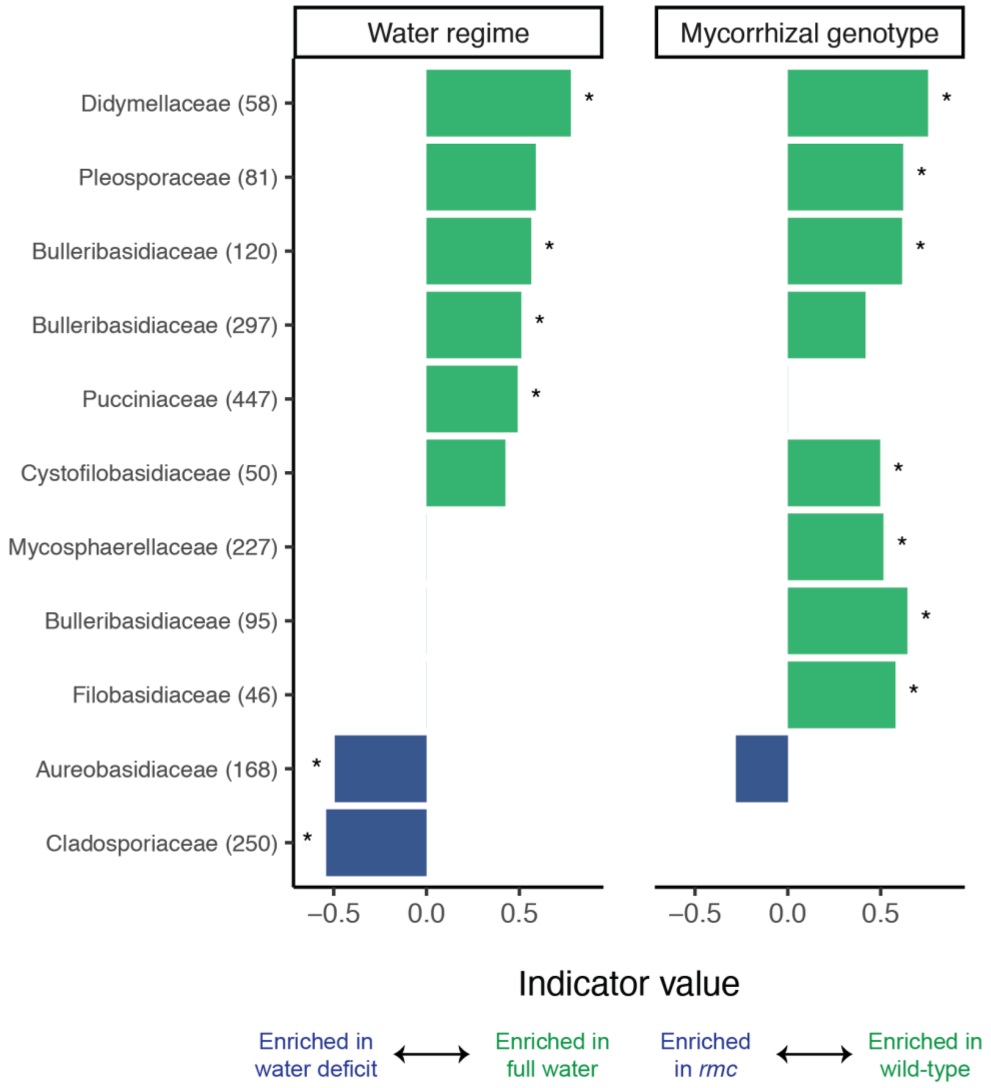

**Figure S8. Overlap in fungal indicator species for water regime and mycorrhizal genotype.** Indicator taxa identified for the water regime (on a wild-type background) or mycorrhizal genotype (on a full water background). Names indicate family-level classification, with ASV number in parenthesis. Values represent the square root of Dufrêne and Legendre's indicator value. For ease of interpretation, the indicator values of opposing treatments (water deficit vs. full water, reduced mycorrhizal vs. wild-type) are displayed with opposing signs. Asterisks indicate nominal significance ($p<0.05$). Missing bars indicate taxa that were too prevalent in both treatments to statistically test association.

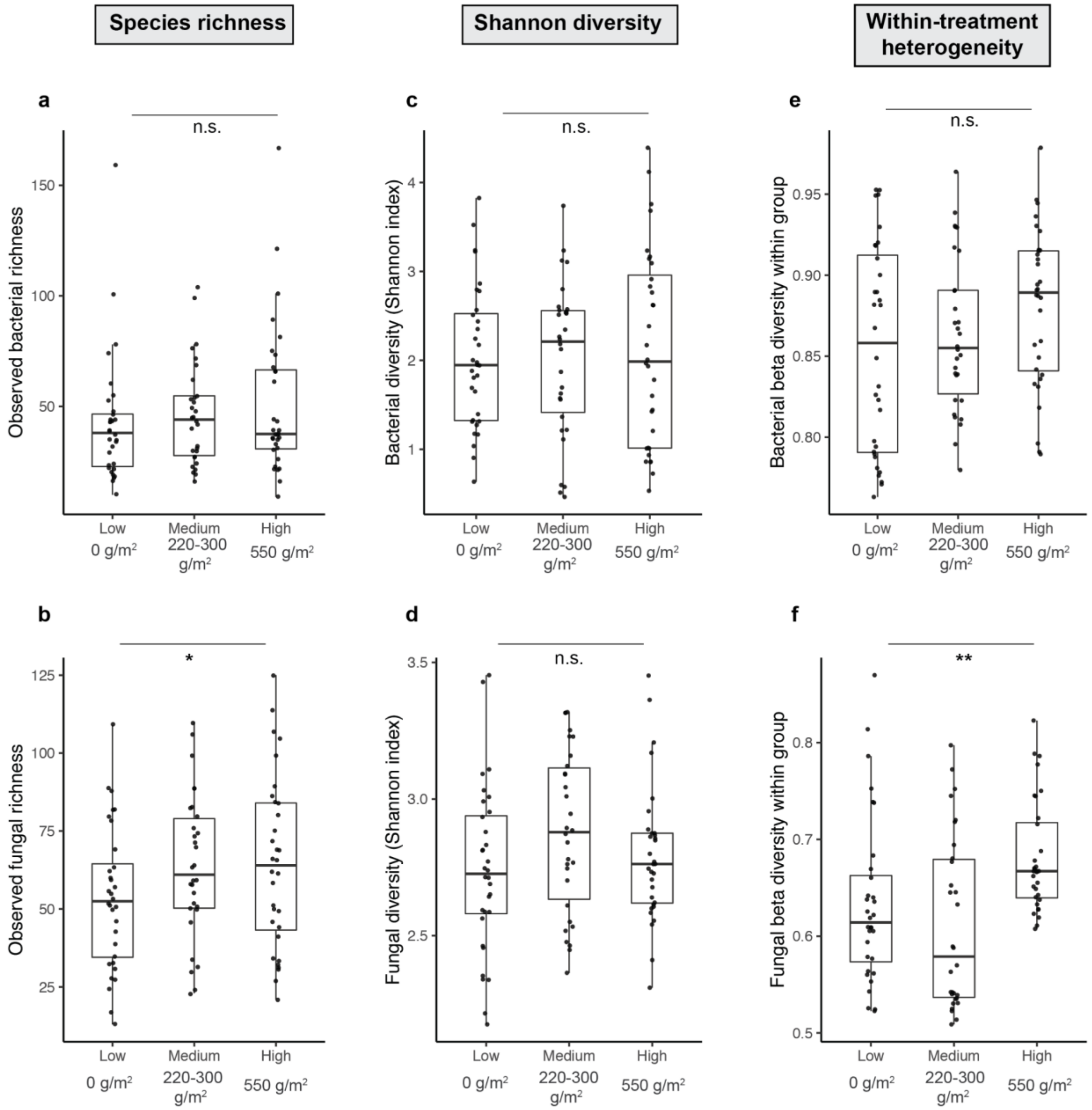

**Figure S9. Phosphorus fertilization increases fungal alpha and beta diversity. (a,b)** ASV counts after removing plant sequences and contaminant sequences. Bacterial richness values were log-transformed prior to ANOVA to meet normality and homoscedasticity assumptions. **(c,d)** Shannon-Weaver index of combined richness and evenness. **(e,f)** Within-group dispersion was quantified by generating a Bray-Curtis dissimilarity matrix. Points represent the mean Bray-Curtis distance of each sample to other samples within the same treatment group (full water or drought).

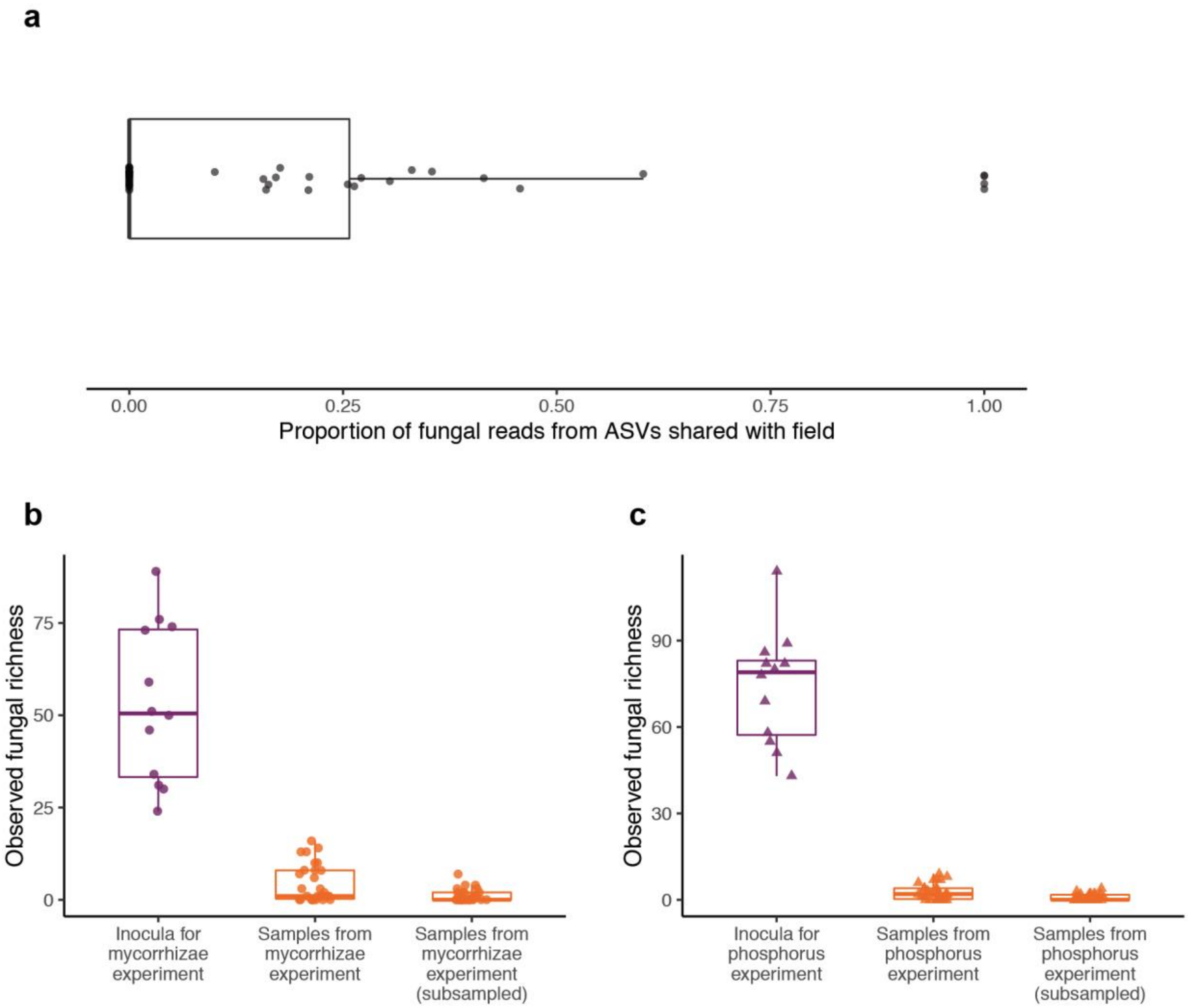

**Figure S10. Colonization of growth chamber plants by field fungal communities. (a)** Fungal communities on plants in the genotype and phosphorus experiments were not dominated by taxa from the field, but rather by taxa unique to the growth chamber. **(b,c)** Removing taxa unique to the growth chamber for further analysis, as was done in the bacterial microbiome, greatly reduces fungal richness values (most plants had only 0-2 fungal ASVs after subsampling).

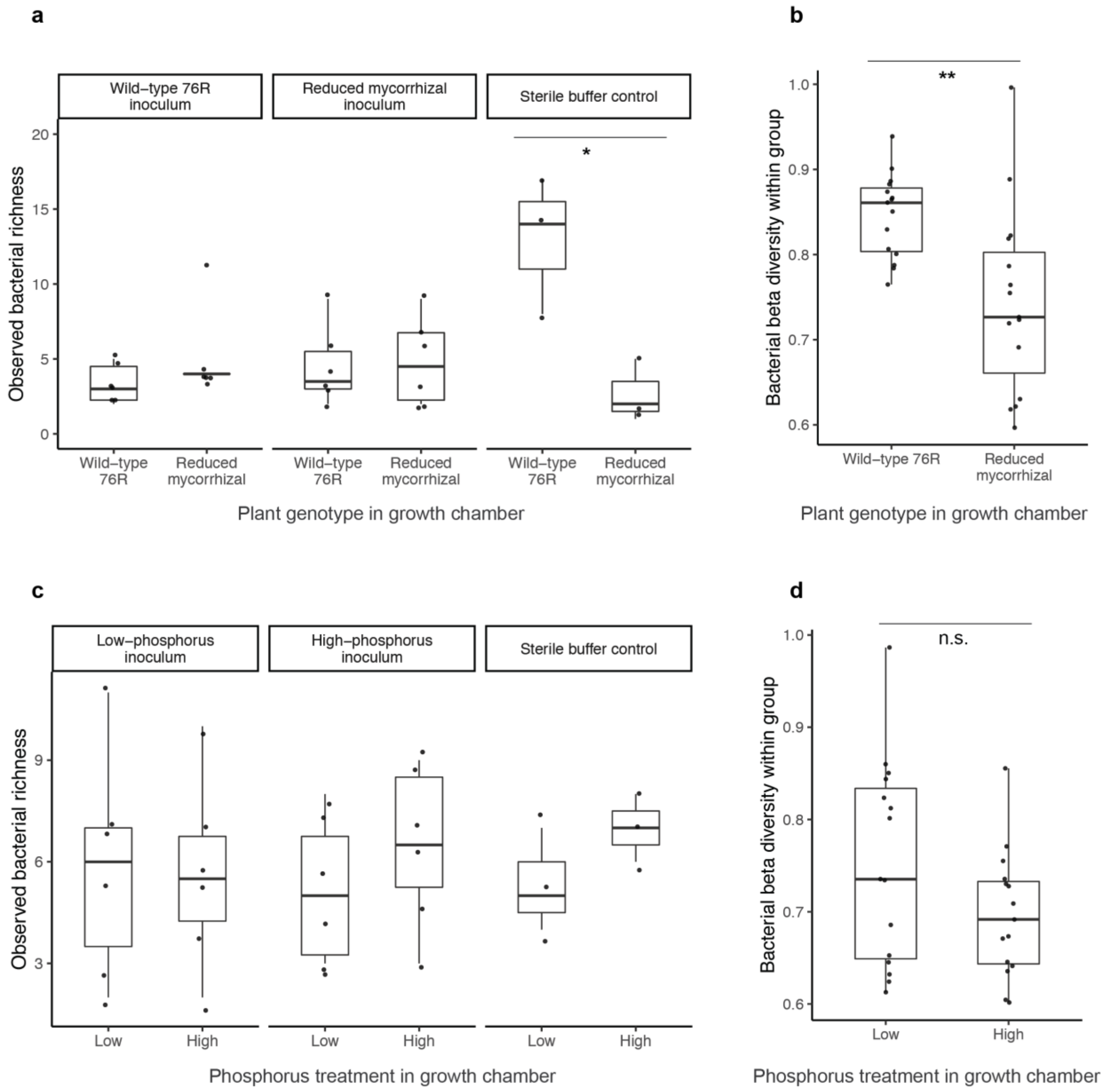

**Figure S11. Growth chamber experiments recapitulate effect of *rmc* genotype, and lack of effect of phosphorus fertilization, on bacterial alpha and beta diversity. (a)** Observed bacterial richness in genotype experiment, as measured by ASV counts after removing chloroplast, mitochondria, and contaminant sequences. Richness values were log-transformed prior to ANOVA to meet normality and homoscedasticity assumptions. **(b)** Within-group dispersion of bacterial communities was quantified by generating a Bray-Curtis dissimilarity matrix. Points represent the mean Bray-Curtis distance of each sample to other samples of the same genotype (76R or reduced mycorrhizal). **(c)** Observed bacterial richness in phosphorus experiment, as measured by ASV counts after removing chloroplast, mitochondria, and contaminant sequences. Richness values were log-transformed prior to ANOVA to meet normality assumption. **(d)** Within-group dispersion of bacterial communities was quantified by generating a Bray-Curtis dissimilarity matrix. Points represent the mean Bray-Curtis distance of each sample to other samples within the same treatment group (low, high).

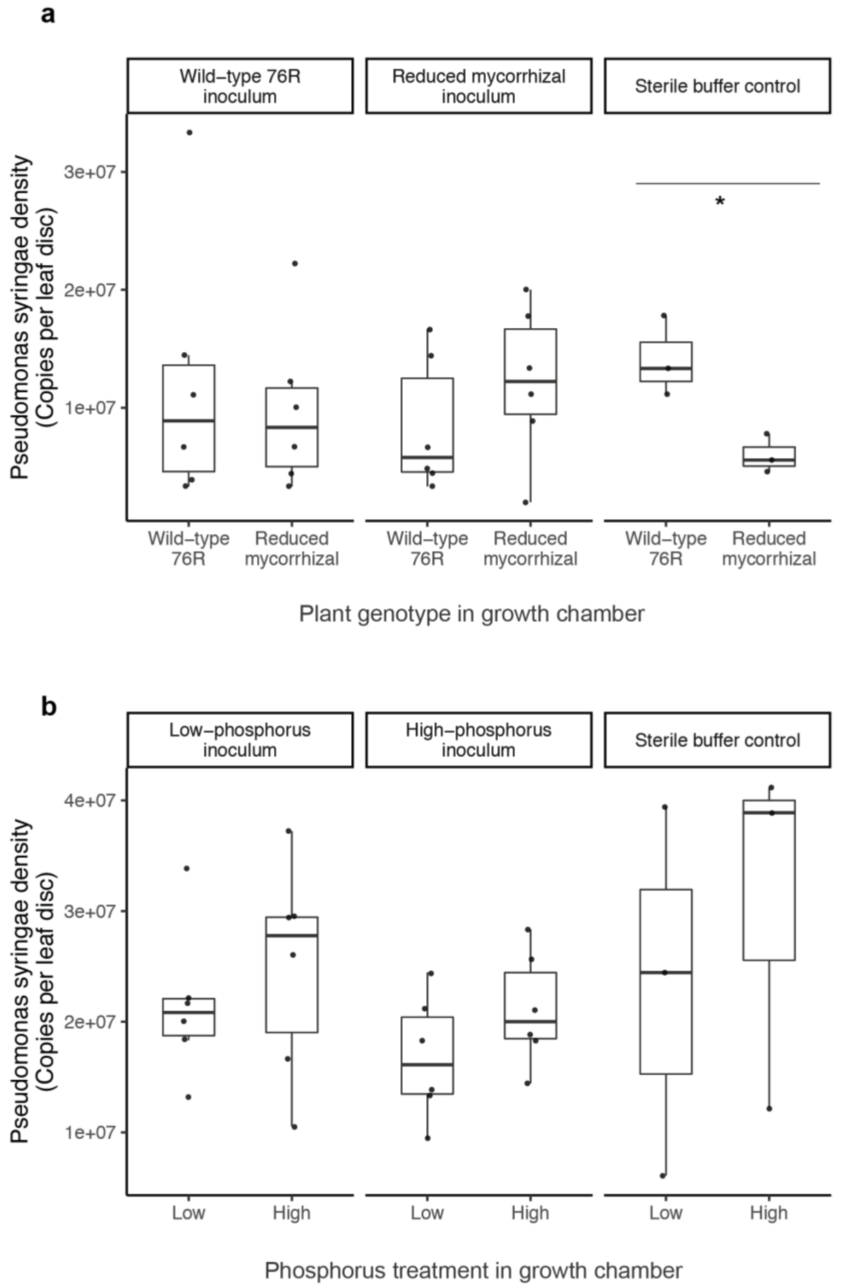

**Figure S12. *Pseudomonas syringae* density on growth chamber plants after experimental infection. (a,b)** Colony forming units (CFUs) on leaf discs (6 mm diameter) clipped from leaves experimentally inoculated with *Pseudomonas syringae* pathovar tomato PT23.

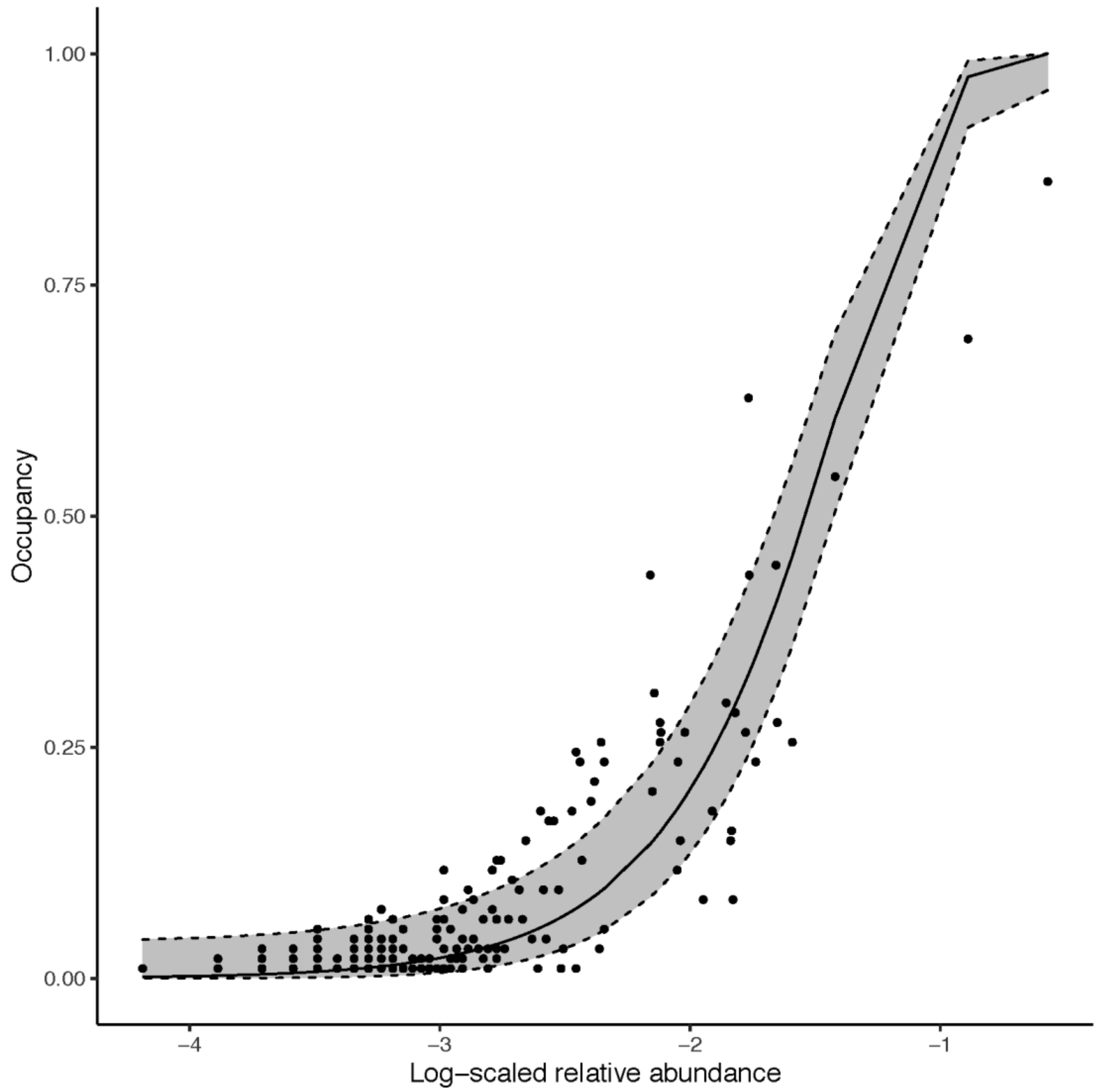

**Figure S13. Occupancy-abundance relationship for bacterial taxa in the field study.** The mean relative abundance of each ASV was plotted against the proportion of plants in which it was detected. The shaded region represents a 95% confidence interval based on the neutral model described by (Sloan et al. 2006) and implemented in the R package *sncm* by (Burns et al. 2016).

Chen, Qiang, William A. Meyer, Qiuwei Zhang, and James F. White. 2020. “16S rRNA Metagenomic Analysis of the Bacterial Community Associated with Turf Grass Seeds from Low Moisture and High Moisture Climates.” *PeerJ* 8 (January): e8417.

Dai, Liangxiang, Guanchu Zhang, Zipeng Yu, Hong Ding, Yang Xu, and Zhimeng Zhang. 2019. “Effect of Drought Stress and Developmental Stages on Microbial Community Structure and Diversity in Peanut Rhizosphere Soil.” *International Journal of Molecular Sciences*. https://doi.org/10.3390/ijms20092265.

Danish, Subhan, Muhammad Zafar-Ul-Hye, Shahid Hussain, Muhammad Riaz, and Muhammad Farooq Qayyum. 2020. “Mitigation of Drought Stress in Maize through Inoculation with Drought Tolerant ACC Deaminase Containing PGPR under Axenic Conditions.” *Pakistan Journal of Botany*. https://doi.org/10.30848/pjb2020-1(7).

Geng, Li-Li, Gao-Xiang Shao, Ben Raymond, Mei-Ling Wang, Xiao-Xiao Sun, Chang-Long Shu, and Jie Zhang. 2018. “Subterranean Infestation by Holotrichia Parallela Larvae Is Associated with Changes in the Peanut (Arachis Hypogaea L.) Rhizosphere Microbiome.” *Microbiological Research* 211 (June): 13–20.

Ghosh, Daipayan, Anshika Gupta, and Sridev Mohapatra. 2019. “A Comparative Analysis of Exopolysaccharide and Phytohormone Secretions by Four Drought-Tolerant Rhizobacterial Strains and Their Impact on Osmotic-Stress Mitigation in Arabidopsis Thaliana.” *World Journal of Microbiology & Biotechnology* 35 (6): 90.

Hone, Holly, Ross Mann, Guodong Yang, Jatinder Kaur, Ian Tannenbaum, Tongda Li, German Spangenberg, and Timothy Sawbridge. n.d. “Drought Tolerant Wheat Varieties Enrich Seed Microbiomes For Beneficial Microbes Under Drought Conditions.” https://doi.org/10.21203/rs.3.rs-113919/v1.

Jang, Seok-Won, Myeong-Hyun Yoou, Woo-Jong Hong, Yeon-Ju Kim, Eun-Jin Lee, and Ki-Hong Jung. 2020. “Re-Analysis of 16S Amplicon Sequencing Data Reveals Soil Microbial Population Shifts in Rice Fields under Drought Condition.” *Rice*. https://doi.org/10.1186/s12284-020-00403-6.

Kaplin, V. G. 2019. “The Main Ways of Infection of Grain Sorghum by Burkholderia Andropogonis (Burkholderiaceae, Burkholderiales) in the Forest-Steppe of the Middle East Volga Region.” *Agrarian Science*. https://doi.org/10.32634/0869-8155-2019-326-1-171-174.

Kavamura, Vanessa Nessner, Suikinai Nobre Santos, João Luiz da Silva, Márcia Maria Parma, Luciana

Aparecida Ávila, Alexandre Visconti, Tiago Domingues Zucchi, Rodrigo Gouvêa Taketani, Fernando Dini Andreote, and Itamar Soares de Melo. 2013. "Screening of Brazilian Cacti Rhizobacteria for Plant Growth Promotion under Drought." *Microbiological Research*. https://doi.org/10.1016/j.micres.2012.12.002.

Kong, Hyun G., Byung K. Kim, Geun C. Song, Soohyun Lee, and Choong-Min Ryu. 2016. "Aboveground Whitefly Infestation-Mediated Reshaping of the Root Microbiota." *Frontiers in Microbiology* 7 (September): 1314.

Li, Hongwei, L. I. Hongwei, Xueling Ding, Chao Wang, K. E. Hongjiao, W. U. Zhou, Yunpeng Wang, L. I. U. Hongxia, and G. U. O. Jianhua. 2016. "Control of Tomato Yellow Leaf Curl Virus Disease by Enterobacter asburiaeBQ9 as a Result of Priming Plant Resistance in Tomatoes." *TURKISH JOURNAL OF BIOLOGY*. https://doi.org/10.3906/biy-1502-12.

Lundberg, Derek S., Scott Yourstone, Piotr Mieczkowski, Corbin D. Jones, and Jeffery L. Dangl. 2013. "Practical Innovations for High-Throughput Amplicon Sequencing." *Nature Methods* 10 (10): 999–1002.

Masyahit, Masanto, Kamaruzaman Sijam, Yahya Awang, and Mohd Ghazali Mohd Satar. 2009. "The First Report of the Occurrence of Anthracnose Disease Caused by Colletotrichum Gloeosporioides (Penz.) Penz. & Sacc. on Dragon Fruit (Hylocereus Spp.) in Peninsular Malaysia." *American Journal of Applied Sciences*. https://doi.org/10.3844/ajas.2009.902.912.

Mathur, Piyush, and Swarnendu Roy. 2021. "Insights into the Plant Responses to Drought and Decoding the Potential of Root Associated Microbiome for Inducing Drought Tolerance." *Physiologia Plantarum*, January. https://doi.org/10.1111/ppl.13338.

Ma, Weihua, Xianyun Zheng, Lixin Li, Jinshan Shen, Wanghong Li, and Ye Gao. 2020. "Changes in the Gut Microbiota of Honey Bees Associated with Jujube Flower Disease." *Ecotoxicology and Environmental Safety*. https://doi.org/10.1016/j.ecoenv.2020.110616.

Meisner, Annelein, Samuel Jacquiod, Basten L. Snoek, Freddy C. Ten Hooven, and Wim H. van der Putten. 2018. "Drought Legacy Effects on the Composition of Soil Fungal and Prokaryote Communities." *Frontiers in Microbiology* 9 (March): 294.

Merenstein, Carter, Jeremy Ward, and David Allen. 2020. "Diplorickettsia Bacteria in an Ixodes Scapularis Tick, Vermont, USA." *Emerging Infectious Diseases*. https://doi.org/10.3201/eid2605.191135.

Naveed, Muhammad, Birgit Mitter, Thomas G. Reichenauer, Krzysztof Wieczorek, and Angela Sessitsch. 2014. "Increased Drought Stress Resilience of Maize through Endophytic Colonization by Burkholderia Phytofirmans PsJN and Enterobacter Sp. FD17." *Environmental and Experimental Botany*. https://doi.org/10.1016/j.envexpbot.2013.09.014.

Nishijima, Kate A. 1987. "Internal Yellowing, a Bacterial Disease of Papaya Fruits Caused by Enterobacter Cloacae." *Plant Disease*. https://doi.org/10.1094/pd-71-1029.

Raheem, Asif, Alexander Shaposhnikov, Andrey A. Belimov, Ian C. Dodd, and Basharat Ali. 2018. "Auxin Production by Rhizobacteria Was Associated with Improved Yield of Wheat (Triticum Aestivum L.) under Drought Stress." *Archives of Agronomy and Soil Science*. https://doi.org/10.1080/03650340.2017.1362105.

Rolli, Eleonora, Ramona Marasco, Gianpiero Vigani, Besma Ettoumi, Francesca Mapelli, Maria Laura Deangelis, Claudio Gandolfi, et al. 2015. "Improved Plant Resistance to Drought Is Promoted by the Root-Associated Microbiome as a Water Stress-Dependent Trait." *Environmental Microbiology* 17 (2): 316–31.

Santos-Medellin, Christian, Zachary Liechty, Joseph Edwards, Bao Nguyen, Bihua Huang, Bart C. Weimer, and Venkatesan Sundaresan. n.d. "Drought Duration Determines the Recovery Dynamics of Rice Root Microbiomes." https://doi.org/10.1101/2020.09.29.314781.

Sloan, William T., Mary Lunn, Stephen Woodcock, Ian M. Head, Sean Nee, and Thomas P. Curtis. 2006. "Quantifying the Roles of Immigration and Chance in Shaping Prokaryote Community Structure." *Environmental Microbiology* 8 (4): 732–40.

Soares, Siomar C., Artur Silva, Eva Trost, Jochen Blom, Rommel Ramos, Adriana Carneiro, Amjad Ali, et al. 2013. "The Pan-Genome of the Animal Pathogen Corynebacterium Pseudotuberculosis Reveals Differences in Genome Plasticity between the Biovar Ovis and Equi Strains." *PloS One* 8 (1): e53818.

Tallapragada, Padmavathi, Rashmi Dikshit, and Swetha Seshagiri. 2016. "Influence of Rhizophagus Spp. andBurkholderia Seminalison the Growth of Tomato (Lycopersicon Esculatum) and Bell Pepper (Capsicum Annuum) under Drought Stress." *Communications in Soil Science and Plant Analysis*.

https://doi.org/10.1080/00103624.2016.1216561.
Verma, Priyanka, Ajar Nath Yadav, Kazy Sufia Khannam, Sanjay Kumar, Anil Kumar Saxena, and Archna Suman. 2016. “Molecular Diversity and Multifarious Plant Growth Promoting Attributes of Bacilli Associated with Wheat (Triticum Aestivum L.) Rhizosphere from Six Diverse Agro-Ecological Zones of India.” *Journal of Basic Microbiology* 56 (1): 44–58.
Wang, Xiaojie, Zhichao Zhang, Zhiqiang Yu, Guofeng Shen, Hefa Cheng, and Shu Tao. 2020. “Composition and Diversity of Soil Microbial Communities in the Alpine Wetland and Alpine Forest Ecosystems on the Tibetan Plateau.” *The Science of the Total Environment* 747 (December): 141358.
Wong, Chi T., Yingqi Xu, Akshari Gupta, James A. Garnett, Steve J. Matthews, and Stephen A. Hare. 2015. “Structural Analysis of Haemoglobin Binding by HpuA from the Neisseriaceae Family.” *Nature Communications* 6 (December): 10172.
Zazharskyi, V. V., P. O. Davydenko, O. M. Kulishenko, I. V. Borovik, A. M. Kabar, and V. V. Brygadyrenko. 2020. “Antibacterial and Fungicidal Effect of Ethanol Extracts from Juniperus Sabina, Chamaecyparis Lawsoniana, Pseudotsuga Menziesii and Cephalotaxus Harringtonia.” *Regulatory Mechanisms in Biosystems*. https://doi.org/10.15421/022015.
Zhang, Rui, Lijuan Chen, Zuirong Niu, Shuzhen Song, and Yan Zhao. 2019. “Water Stress Affects the Frequency of Firmicutes, Clostridiales and Lysobacter in Rhizosphere Soils of Greenhouse Grape.” *Agricultural Water Management*. https://doi.org/10.1016/j.agwat.2019.105776.
